# TMEM65 regulates NCLX-dependent mitochondrial calcium efflux

**DOI:** 10.1101/2023.10.06.561062

**Authors:** Joanne F. Garbincius, Oniel Salik, Henry M. Cohen, Carmen Choya-Foces, Adam S. Mangold, Angelina D. Makhoul, Anna E. Schmidt, Dima Y. Khalil, Joshua J. Doolittle, Anya S. Wilkinson, Emma K. Murray, Michael P. Lazaropoulos, Alycia N. Hildebrand, Dhanendra Tomar, John W. Elrod

## Abstract

The balance between mitochondrial calcium (_m_Ca^2+^) uptake and efflux regulates ATP production, but if perturbed causes energy starvation or _m_Ca^2+^ overload and cell death. The mitochondrial sodium-calcium exchanger, NCLX, is a critical route of _m_Ca^2+^ efflux in excitable tissues, such as the heart and brain, and animal models support NCLX as a promising therapeutic target to limit pathogenic _m_Ca^2+^ overload. However, the mechanisms that regulate NCLX activity remain largely unknown. We used proximity biotinylation proteomic screening to identify the NCLX interactome and define novel regulators of NCLX function. Here, we discover the mitochondrial inner membrane protein, TMEM65, as an NCLX-proximal protein that potently enhances sodium (Na^+^)-dependent _m_Ca^2+^ efflux. Mechanistically, acute pharmacologic NCLX inhibition or genetic deletion of NCLX ablates the TMEM65-dependent increase in _m_Ca^2+^ efflux. Further, loss-of-function studies show that TMEM65 is required for Na^+^-dependent _m_Ca^2+^ efflux. Co-fractionation and *in silico* structural modeling of TMEM65 and NCLX suggest these two proteins exist in a common macromolecular complex in which TMEM65 directly stimulates NCLX function. In line with these findings, knockdown of *Tmem65* in mice promotes _m_Ca^2+^ overload in the heart and skeletal muscle and impairs both cardiac and neuromuscular function. We further demonstrate that *TMEM65* deletion causes excessive mitochondrial permeability transition, whereas TMEM65 overexpression protects against necrotic cell death during cellular Ca^2+^ stress. Collectively, our results show that loss of TMEM65 function in excitable tissue disrupts NCLX-dependent _m_Ca^2+^ efflux, causing pathogenic _m_Ca^2+^ overload, cell death and organ-level dysfunction, and that gain of TMEM65 function mitigates these effects. These findings demonstrate the essential role of TMEM65 in regulating NCLX-dependent _m_Ca^2+^ efflux and suggest modulation of TMEM65 as a novel strategy for the therapeutic control of _m_Ca^2+^ homeostasis.

---

The mitochondrial sodium-calcium exchanger, NCLX, mediates the extrusion of calcium (Ca^2+^) from the mitochondrial matrix in exchange for the entry of sodium (Na^+^) and is the primary route for mitochondrial Ca^2+^ (_m_Ca^2+^) efflux in excitable cells^1,2^. We recently demonstrated the requirement of NCLX in maintaining _m_Ca^2+^ homeostasis and preventing pathogenic _m_Ca^2+^ overload in cardiomyocytes^3^ and neurons^4^. Moreover, we and others have demonstrated the efficacy of augmenting NCLX expression and activity to limit disease progression in models of ischemia-reperfusion (I/R) injury and ischemic heart failure^3^, non-ischemic heart failure^5^, Alzheimer’s disease^6^, and cancer^7^. These results support the modulation of NCLX activity as a promising therapeutic strategy for a wide range of diseases. However, a critical barrier to clinical translation is a lack of therapeutic modulators of NCLX activity, largely due to our very limited understanding of the mechanisms regulating NCLX function. Knowing that the activity of plasma membrane Na^+^/Ca^2+^ exchangers that are structurally related to NCLX are regulated by allosteric interactions with other proteins^8–10^, we hypothesized that mapping the NCLX protein interactome would reveal regulators of NCLX function and provide new targets to modulate _m_Ca^2+^ efflux^11^. Here, we report the identification of TMEM65, an inner mitochondrial membrane protein, as the first genetically-confirmed regulator of NCLX-dependent _m_Ca^2+^ efflux and demonstrate its critical role in preventing pathogenic _m_Ca^2+^ overload in excitable tissues.

## Discovery of TMEM65 as a mitochondrial localized NCLX interactor

Despite growing support for NCLX as a therapeutic target in diseases featuring _m_Ca^2+^ overload, progress towards defining the regulation of NCLX activity has been stymied by numerous technical challenges that complicate the study of this protein. NCLX has a complex structure comprising 13 hydrophobic transmembrane domains^12^; it is embedded in the highly complex lipid environment of the inner mitochondrial membrane^2^; no highly specific pharmacologic modulators of its function exist^11^; and the questionable validity and inconsistent commercial availability of antibodies directed against it^7,11,13^ are major hurdles for the field. Therefore, we generated a fusion construct encoding human NCLX protein followed by a biotin ligase domain (BioID2) and C-terminal HA-tag (Fig. 1a) to interrogate the NCLX interactome. Expression of NCLX-BioID2-HA in intact cells and supplementation with biotin allows for biotinylation of lysine residues in proteins that come within a vicinity of <10nm^14,15^. Cellular fractionation of human AC16 cardiomyocytes transiently transfected with NCLX-BioID2-HA confirmed its localization to mitochondria (Fig. 1b). Progressive digestion of mitochondria from AC16 cardiomyocytes transiently expressing this construct with increasing concentrations of trypsin indicated that despite increasing degradation of the full-length fusion protein, the ∼25kD C-terminal BioID2-HA moiety was protected from proteolysis (Extended Data Fig.1). After comparing the NCLX- BioID2-HA degradation pattern to the degradation of the outer mitochondrial membrane protein TOM20, the inner mitochondrial membrane (IMM) protein MCU, and the mitochondrial matrix protein Cyclophilin D (Extended Data Fig. 1), we interpreted that NCLX-BioID2-HA properly localized to the IMM with the C-terminal tail positioned in the mitochondrial matrix (Fig. 1a). Blotting with streptavidin revealed the persistence of biotinylated proteins even at the highest concentration of trypsin (Extended Data Fig. 1), further supporting proper IMM localization and orientation of the NCLX-BioID2-HA fusion protein. Biotinylated proteins from whole cell lysates of AC16 cardiomyocytes transiently expressing NCLX-BioID2-HA, or control BioID2-HA alone (Fig. 1c), were identified by mass spectrometry. Biotinylated proteins enriched ≥2-fold with NCLX-BioID2-HA compared to BioID2-HA in a minimum of 2 out of 3 replicate experiments were cross-referenced to the Integrated Mitochondrial Protein Index (IMPI) to identify NCLX- proximal mitochondrial proteins (Extended Data Table 1). This analysis identified transmembrane protein 65 (TMEM65), an IMM protein of unknown molecular function^16^, as a potential NCLX-interactor in cardiomyocytes (Fig. 1d).

**Fig. 1:**
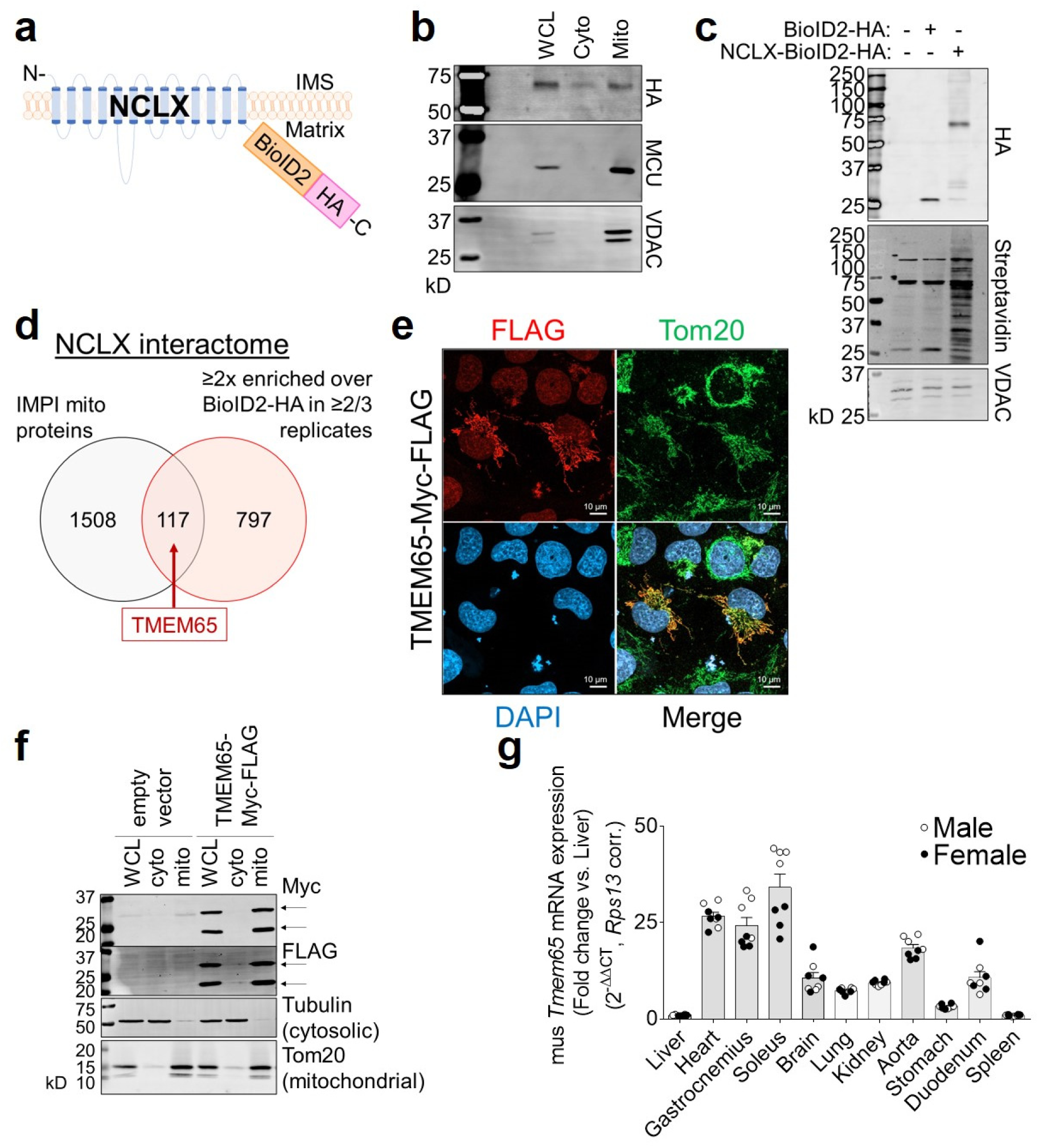
*In vitro* proximity biotinylation screen identifies TMEM65 as a member of the NCLX interactome. a, Schematic of human NCLX-BioID2-HA fusion protein. IMS, intermembrane space. b, Western blotting of cellular fractions of AC16 cardiomyocytes transiently transfected with a plasmid encoding NCLX-BioID2-HA. MCU and VDAC served as mitochondrial markers. WCL, whole cell lysate; Cyto, cytosolic fraction, Mito, mitochondrial fraction. c, Expression of BioID2-HA and NCLX-BioID2-HA fusion protein and resulting cellular biotinylation. d, Summary of mass spectrometry results from proximity biotinylation screen in AC16 cardiomyocytes. Overlapping region of Venn diagram represents known or predicted mitochondrial proteins according to the IMPI, the biotinylation of which were enriched in NCLX-BioID2-HA samples compared to BioID2-HA negative controls. e, Immunofluorescence staining of AC16 cardiomyocytes transiently transfected with TMEM65-Myc-FLAG. Tom20 served as a mitochondrial marker. f, Western blots showing cellular fractionation and mitochondrial localization of exogenously expressed TMEM65-Myc-FLAG protein (arrows) in AC16 cardiomyocytes. g, *Tmem65* mRNA expression in adult male (open symbols) and female (filled symbols) mouse tissues (*n* = 8 mice).

**Extended Data Fig. 1:**
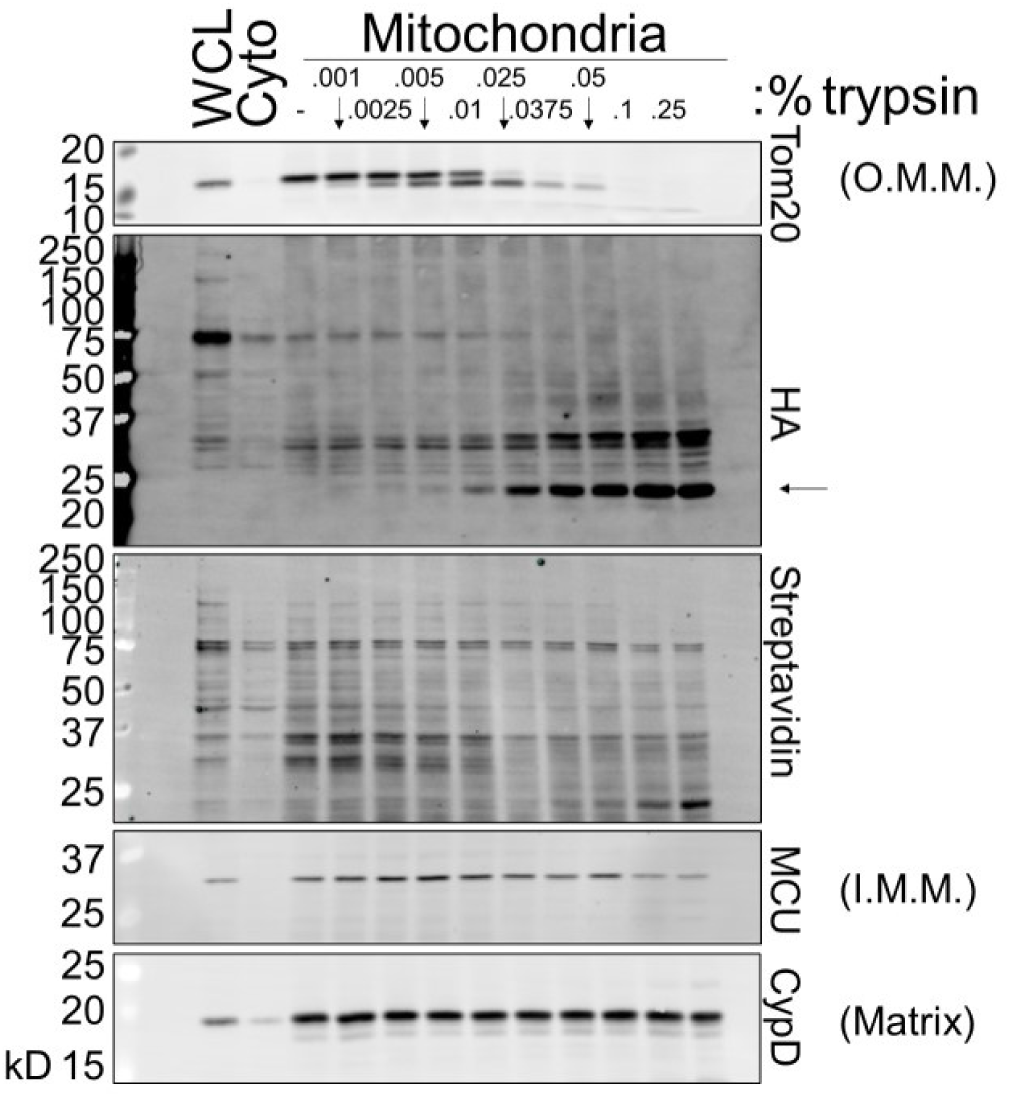
Validation of mitochondrial localization of NCLX-BioID2-HA fusion protein. Western blots showing progressive digestion of mitochondria from AC16 cardiomyocytes expressing NCLX-BioID2HA with increasing concentrations of trypsin. Arrow indicates ∼25-kD BioID2-HA moiety. WCL, whole cell lysate; Cyto, cytosolic fraction. CypD, cyclophilin D. O.M.M., outer mitochondrial membrane; I.M.M., inner mitochondrial membrane.

We prioritized TMEM65 for further analysis given a recent human case report of mitochondrial encephalomyopathy in a patient with a homozygous null mutation in *TMEM65*^17^ and the resemblance of the resulting mitochondrial and organ-level dysfunction to that which is observed in human patients and mouse models with genetic mutations that perturb mitochondrial calcium homeostasis^18–21^. Immunofluorescence staining (Fig. 1e) and cellular fractionation (Fig. 1f) confirmed mitochondrial localization of exogenous Myc-FLAG-tagged TMEM65 in AC16 cardiomyocytes. Fitting with the detection of TMEM65 as part of the NCLX interactome in AC16 cardiomyocytes, qPCR revealed murine *Tmem65* gene expression to be enriched in striated muscle and other excitable tissues, including the brain (Fig. 1g).

## TMEM65 enhances and is required for Na^+^-dependent _m_Ca^2+^ efflux

We next tested the effect of TMEM65 overexpression (OE) on _m_Ca^2+^ flux *in vitro*. We generated clonal lines of AC16 cardiomyocytes with stable expression of C-terminal epitope-tagged human TMEM65 (TMEM65-Myc-FLAG) (Fig. 2a, Extended Data Fig. 2a). These cells exhibited a ∼65% reduction in NCLX protein expression (Fig. 2a, Extended Data Fig. 2b), consistent with the notion that TMEM65 and NCLX have a shared effect on _m_Ca^2+^ handling. Stable TMEM65-OE increased the inhibitory phosphorylation of pyruvate dehydrogenase (PDH) Ser293 (Extended Data Fig. 2c-d), indicating diminished steady-state matrix-Ca^2+^ content. In permeabilized AC16 cardiomyocytes incubated with thapsigargin to inhibit sarco/endoplasmic reticulum Ca^2+^ uptake, stable TMEM65-OE enhanced the rate of _m_Ca^2+^ efflux (Fig. 2b-c), as measured following acute inhibition of _m_Ca^2+^ uptake with the mitochondrial calcium uniporter inhibitor, Ru360. TMEM65- OE also attenuated the initial _m_Ca^2+^ uptake rate (Extended Data Fig. 2e) and net _m_Ca^2+^ uptake prior to the addition of Ru360 (Fig. 2d), consistent with previous reports describing that _m_Ca^2+^ uptake and efflux are ongoing processes that occur simultaneously^3^. To verify that these effects were not attributable to secondary compensatory effects of stable TMEM65 overexpression, we also assessed _m_Ca^2+^ flux in permeabilized AC16 cells after acute adenoviral TMEM65 transduction (Extended Data Fig. 2f). Consistent with our findings in stable cell lines, acute TMEM65 overexpression enhanced the rate and extent of _m_Ca^2+^ efflux (Extended Data Fig. 2g-i), and limited the extent of acute net _m_Ca^2+^ uptake (Extended Data Fig. 2j).

**Fig. 2:**
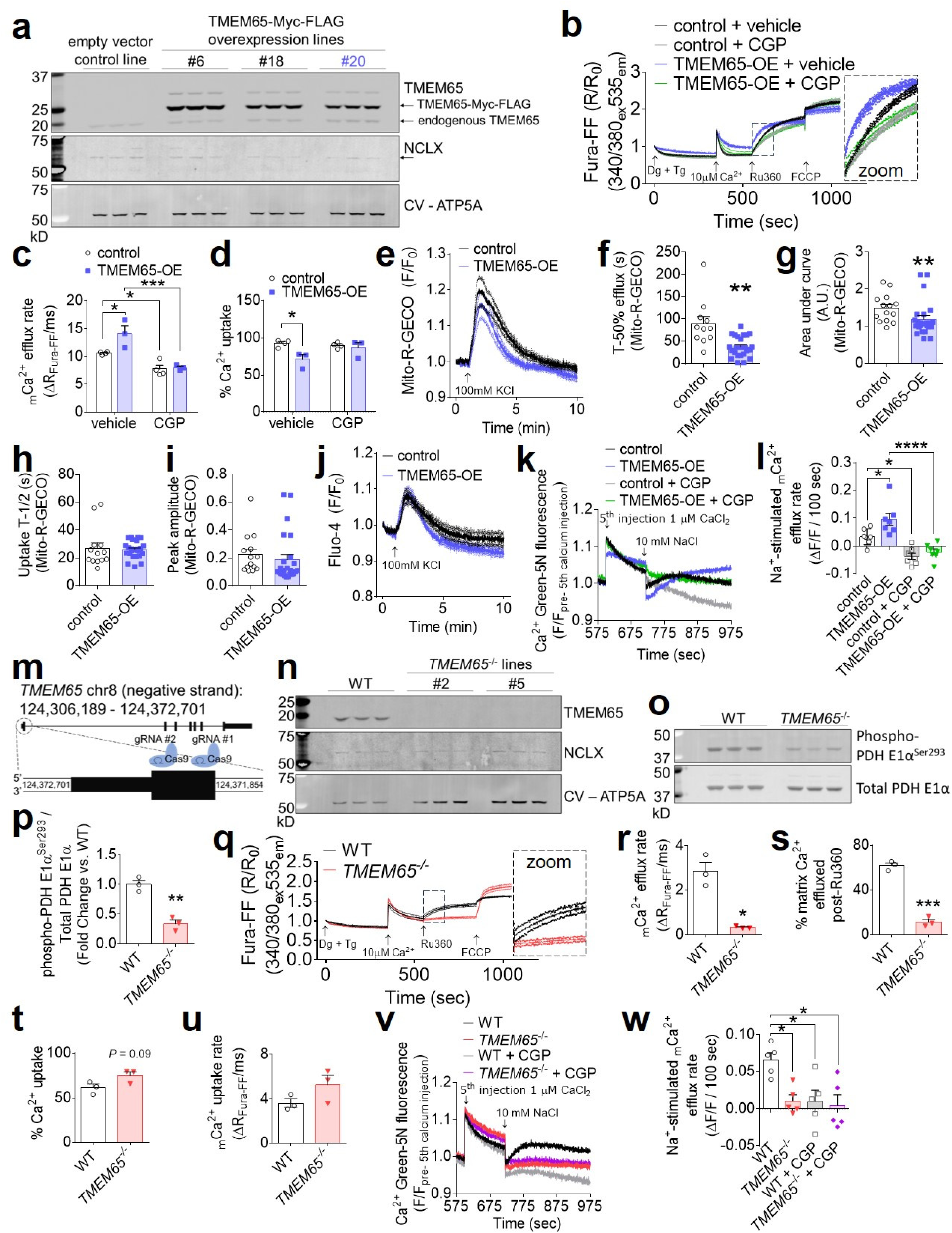
Gain of TMEM65 function enhances and loss of TMEM65 impairs sodium-dependent mitochondrial calcium efflux. a, Western blotting of AC16 cardiomyocytes stably transduced with control empty vector or TMEM65-Myc-FLAG. Corresponding densitometric quantifications are shown in Extended Data Fig. 2 a-b. Line #20 (blue) was used in subsequent experiments. b, Pooled traces for AC16 cardiomyocytes with stable overexpression of TMEM65- Myc-FLAG (TMEM65-OE) or empty vector control, in the presence or absence of the NCLX inhibitor CGP-37157 (CGP), representing extramitochondrial bath Ca^2+^ (Fura-FF). 2 million cells were permeabilized with digitonin (Dg) in the presence of thapsigargin (Tg), then administered a single 10µM Ca^2+^ bolus, with subsequent additions of the mitochondrial calcium uniporter inhibitor Ru360 and FCCP as indicated. (*n* = 4 for control, 3 for TMEM65-OE). Inset shows initial _m_Ca^2+^ efflux phase. _m_Ca^2+^ efflux rate over the first 25 seconds following addition of Ru360 (c) and % net _m_Ca^2+^ uptake following the 10µM Ca^2+^ bolus (d). (*n* = 4 for control, 3 for TMEM65- OE. \**P*< 0.05, \*\**P*<0.01, \*\*\**P*< 0.001, by one-way ANOVA with Sidak’s post-hoc test). e, Pooled traces showing KCl-evoked mitochondrial Ca^2+^ transients as measured by Mito-R-GECO fluorescence in intact AC16 cardiomyocytes. (*n* = 14 cells for empty vector control, 24 cells for TMEM65-OE). f, Time to 50% efflux for _m_Ca^2+^ transients (*n* = 11 cells for empty vector control, 24 cells for TMEM65-OE; \*\**P*< 0.01 by unpaired t-test with Welch’s correction). g, Area under the curve for _m_Ca^2+^ transients (*n* = 14 cells for empty vector control, 24 cells for TMEM65-OE; \*\**P*< 0.01 by Mann-Whitney test). h, Half rise time of _m_Ca^2+^ transients (*n* = 13 cells for empty vector control, 24 cells for TMEM65-OE, n.s. by Mann-Whitney test). i, Peak amplitude of _m_Ca^2+^ transients. (*n* = 14 cells for empty vector control, 24 cells for TMEM65-OE, n.s. by Mann-Whitney test). j, Pooled traces showing KCl-evoked cytosolic Ca^2+^ transients as measured by Fluo-4 fluorescence in intact AC16 cardiomyocytes. (*n* = 9 cells/group). k, Mean of pooled traces for plate reader _m_Ca^2+^ flux assays with permeabilized AC16 cells. Traces show extramitochondrial Ca^2+^ (Ca^2+^ Green 5N fluorescence), beginning with addition of the final of 5 consecutive 1-µM Ca^2+^ boluses, followed by stimulation of _m_Ca^2+^ efflux by the addition of 10mM NaCl. GCP, CGP-37157. Error bars were omitted for clarity. (*n* = 7/group). l, Quantification of Na^+^-stimulated and CGP-sensitive _m_Ca^2+^ efflux (n = 7/group, \**P*<0.05, \*\*\*\**P*<0.001 by 1-way ANOVA with Tukey’s post-hoc test). m, Approach for CRISPR/Cas9-mediated disruption of *TMEM65*. n, Western blot confirming loss of TMEM65 protein expression in clonal AC16 cardiomyocyte lines following CRISPR/Cas9-mediated gene editing. o, Western blots for pyruvate dehydrogenase (PDH) phosphorylation status in wild-type (WT) and *TMEM65*^-/-^ AC16 cardiomyocytes. p, Densitometric quantification of PDH phosphorylation normalized to total PDH. (*n* = 3/group; \*\**P*<0.01 by unpaired t-test). q, Pooled traces for WT and *TMEM65*^-/-^ AC16 cardiomyocytes showing extramitochondrial bath Ca^2+^ (Fura-FF). 4 million cells were permeabilized with digitonin (Dg) in the presence of thapsigargin (Tg), then administered a single 10µM Ca^2+^ bolus, with subsequent additions of the mitochondrial calcium uniporter inhibitor Ru360 and FCCP as indicated. (*n* = 3/group). r, _m_Ca^2+^ efflux rate over the first 25 seconds following addition of Ru360. (*n* = 3/group, \**P*<0.05 by unpaired t-test with Welch’s correction). Percent of matrix Ca^2+^ effluxed post-Ru360 (s); % net _m_Ca^2+^ uptake following the 10µM Ca^2+^ bolus (t); and _m_Ca^2+^ uptake rate over the first 25 seconds following the Ca^2+^ peak (u). (*n* = 3 group. \*\*\**P*< 0.001 by unpaired t-test). v, Mean of pooled traces for plate reader _m_Ca^2+^ flux assays with permeabilized AC16 cells as in (k). Error bars were omitted for clarity. (*n* = 5/group). w, Quantification of Na^+^-stimulated and CGP-sensitive _m_Ca^2+^ efflux (*n* = 5/group, \**P*<0.05 by 1-way ANOVA with Tukey’s post-hoc test).

**Extended Data Fig. 2:**
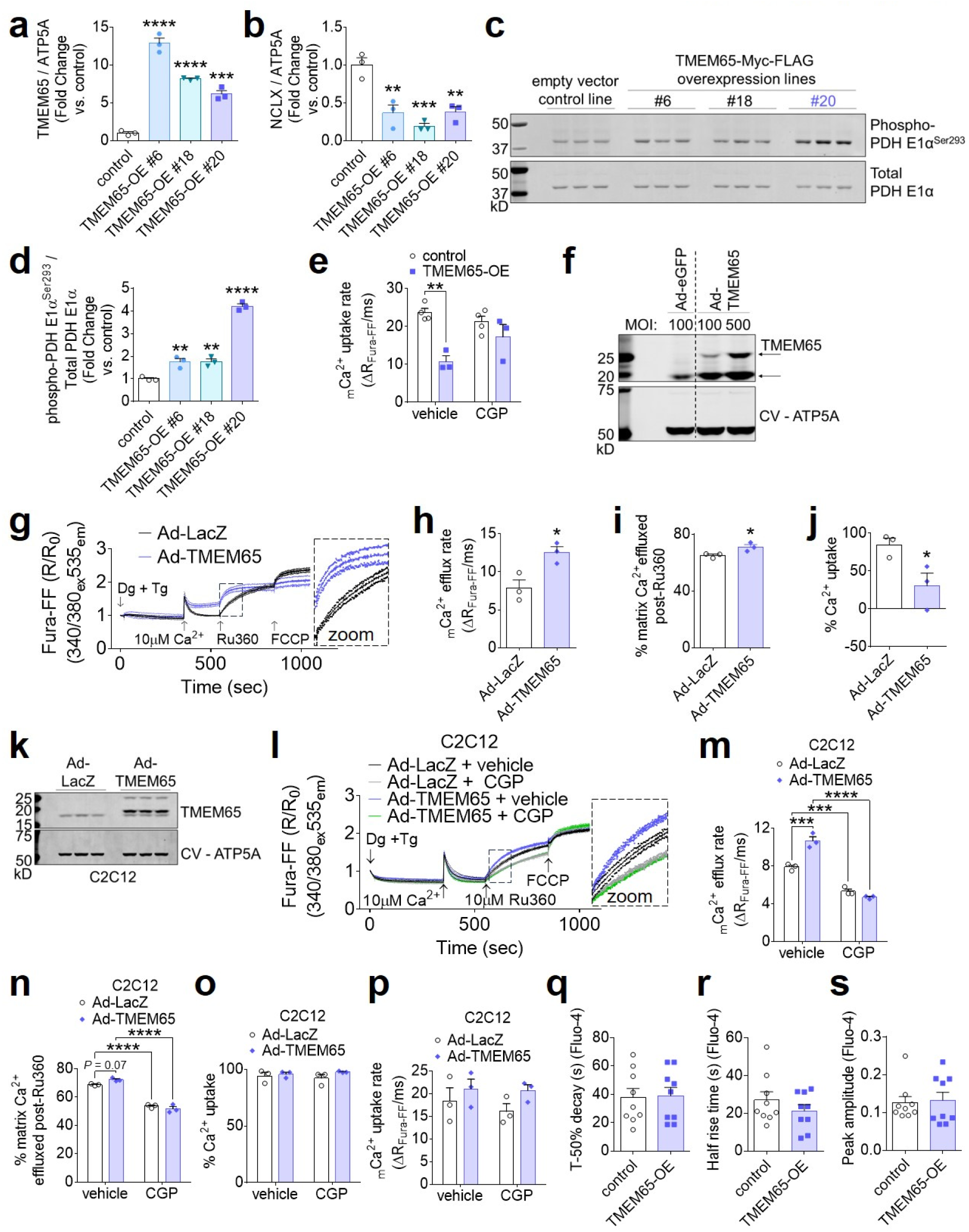
Effects of TMEM65 overexpression on _m_Ca^2+^ and cytosolic Ca^2+^ handling. a-b, Densitometric quantifications of western blots shown in Fig. 2a. c, Western blots for pyruvate dehydrogenase (PDH) phosphorylation status in control and stable TMEM65-Myc-FLAG overexpression lines (TMEM65-OE). d, Densitometric quantification of PDH phosphorylation normalized to total PDH. (*n* = 3/group; \*\**P*<0.01, \*\*\*\**P*<0.0001 by one-way ANOVA with Dunnett’s post-hoc test). e, _m_Ca^2+^ uptake rate for permeabilized cell experiments shown in Fig. 2b. (*n* = 4 for empty vector control, 3 for TMEM65-OE. \*\**P*<0.01 by one-way ANOVA with Sidak’s post-hoc test). f, Western blotting of AC16 cardiomyocytes transduced for 24 hours with adenovirus encoding human TMEM65. MOI, multiplicity of infection. Dashed line is to aid in visualizing separate conditions. g, Pooled traces for AC16 cardiomyocytes transduced with adenovirus encoding LacZ (Ad-LacZ) or human TMEM65 (Ad-TMEM65), showing extramitochondrial bath Ca^2+^ (Fura-FF). 4 million cells were permeabilized with digitonin (Dg) in the presence of thapsigargin (Tg), then administered a single 10µM Ca^2+^ bolus, with subsequent additions of the mitochondrial calcium uniporter inhibitor Ru360 and FCCP as indicated. (*n* = 3/group). Inset shows initial _m_Ca^2+^ efflux phase. _m_Ca^2+^ efflux rate over the first 25 seconds following addition of Ru360 (h); % of matrix Ca^2+^ effluxed post-Ru360 (i); and % net _m_Ca^2+^ uptake following 10µM Ca^2+^ bolus (j). (*n* = 3/genotype; \**P*< 0.05 by unpaired t-test). k, Western blotting of C2C12 myoblasts transduced for 24 hours with adenovirus encoding human TMEM65. l, Pooled traces for 2million mouse C2C12 skeletal myoblasts with acute transduction of Ad-LacZ or Ad-TMEM65, in the presence or absence of the NCLX inhibitor CGP-37157 (CGP), showing extramitochondrial bath Ca^2+^ (Fura-FF) as in (g) (*n* = 3/group). _m_Ca^2+^ efflux rate over the first 25 seconds following addition of Ru360 (m), % of matrix Ca^2+^ effluxed post-Ru360 (n); % net _m_Ca^2+^ uptake following the 10µM Ca^2+^ bolus (o), and initial _m_Ca^2+^ uptake rate for the first 25 seconds following the Ca^2+^ peak (p). (*n* = 3/group. \*\*\**P*< 0.001, \*\*\*\**P*<0.0001 vs. WT by two-way ANOVA with Sidak’s post-hoc test. All comparisons n.s. for panels o and p). Time to 50% decay (q) and half rise time (r) of KCl-evoked cytosolic Ca^2+^ transients as measured by Fluo-4 fluorescence in intact AC16 cardiomyocytes for experiment shown in Fig. 2j. (*n*= 9 cells/group. n.s. in all cases by unpaired t-test). s, peak amplitude of KCl-evoked cytosolic Ca^2+^ transients for experiment shown in Fig. 2j. (*n* = 9 cells/group. n.s. by Mann-Whitney test).

Of note, acute treatment with the pharmacologic NCLX inhibitor, CGP-37157, completely attenuated the TMEM65-dependent increase in _m_Ca^2+^ efflux rate, reduction in net _m_Ca^2+^ uptake and reduction in _m_Ca^2+^ uptake rate in AC16 cardiomyocytes (Fig. 2b-d; Extended Data Fig. 2e). This phenomenon whereby NCLX inhibition abrogated TMEM65’s enhancement of _m_Ca^2+^ efflux was conserved in other striated muscle cell types, including mouse C2C12 skeletal myoblasts (Extended Data Fig. 2k-p). These findings indicate that TMEM65 functionally depends on NCLX activity to augment _m_Ca^2+^ extrusion, and support the hypothesis that TMEM65 functions as a positive regulator of NCLX activity.

Assessment of KCl-evoked _m_Ca^2+^ transients in intact AC16 cardiomyocytes likewise showed that stable TMEM65-OE accelerated _m_Ca^2+^ efflux and reduced net _m_Ca^2+^ accumulation (Fig. 2e- g), without altering _m_Ca^2+^ uptake (Fig. 2h-i). Importantly, no changes in KCl-induced cytosolic Ca^2+^ transients were noted (Fig. 2j, Extended Data Fig. 2q-s). Finally, to further explore the potential NCLX-dependence of TMEM65-sensitive _m_Ca^2+^ efflux, we evaluated Na^+^-induced _m_Ca^2+^ efflux in permeabilized cells. TMEM65-OE specifically accelerated _m_Ca^2+^ efflux stimulated by the acute addition of Na^+^, and this effect was sensitive to pharmacologic NCLX inhibition (Fig. 2k-l). Together, these observations indicate that TMEM65 promotes Na^+^-sensitive _m_Ca^2+^ efflux, likely through the only known mitochondrial Na^+^/Ca^2+^ exchanger, NCLX.

To determine the requirement for TMEM65 in mitochondrial calcium efflux, we used CRISPR/Cas9 to disrupt the *TMEM65* gene in AC16 cardiomyocytes (Fig. 2m). Western blotting confirmed loss of TMEM65 protein in clonal *TMEM65*^-/-^ lines (Fig. 2n). Loss of TMEM65 decreased inhibitory phosphorylation of PDH Ser293 (Fig. 2o-p), suggesting elevated matrix-Ca^2+^ content. Examination of _m_Ca^2+^ exchange in permeabilized cells treated with thapsigargin showed that deletion of *TMEM65* attenuated _m_Ca^2+^ efflux (Fig. 2q-s). The disruption of _m_Ca^2+^ efflux with loss of TMEM65 tended to enhance _m_Ca^2+^ uptake (Fig. 2t-u), further emphasizing that ongoing _m_Ca^2+^ efflux occurs even during acute _m_Ca^2+^ uptake, as we have previously reported^3^. Finally, consistent with our observations in cells with TMEM65 overexpression, loss of TMEM65 specifically abrogated Na^+^-induced _m_Ca^2+^ efflux (Fig. 2v-w). Moreover, GCP-37157-sensitive _m_Ca^2+^ efflux was completely lost in the absence of TMEM65 (Fig 2v-w). These data, our identification of TMEM65 in the NCLX interactome, and the resemblance of the *in vitro* phenotypes resulting from gain-or loss of TMEM65 function to those resulting from gain-or loss of NCLX function^3^, support the hypothesis that TMEM65 enhances _m_Ca^2+^ efflux in an NCLX-dependent manner.

## TMEM65 enhances _m_Ca^2+^ efflux through NCLX

To more rigorously examine the functional interactions between TMEM65 and NCLX in mediating _m_Ca^2+^ efflux, we next evaluated _m_Ca^2+^ transients in immortalized *Slc8b1*^fl/fl^ mouse embryonic fibroblasts (MEFs) following acute deletion of NCLX by adenoviral delivery of Cre recombinase (note that *Slc8b1* encodes NCLX, and that *Slc8b1*^fl/fl^ hereafter is referred to as *Nclx*^fl/fl^). Adenoviral TMEM65 expression accelerated the initial decay of the _m_Ca^2+^ transient (efflux rate) in control *Nclx*^fl/fl^ cells, but not in *Nclx*^-/-^ cells (Fig. 3a-b). We also observed a reduction in the area-under-the-curve of the _m_Ca^2+^ transient with Ad-TMEM65 in control cells, and this effect was lost in *Nclx*^-/-^ cells (Fig. 3c). This indicates that TMEM65 can limit net matrix Ca^2+^ accumulation in response to increased cytosolic Ca^2+^, but that this effect requires NCLX. Note that the change in efflux with Ad-TMEM65 was more modest in these experiments, likely due to more limited Na^+^-dependent _m_Ca^2+^ efflux in fibroblasts in contrast to excitable cells. Yet, these data remain consistent with TMEM65 enhancing _m_Ca^2+^ efflux specifically through effects on NCLX.

**Fig. 3:**
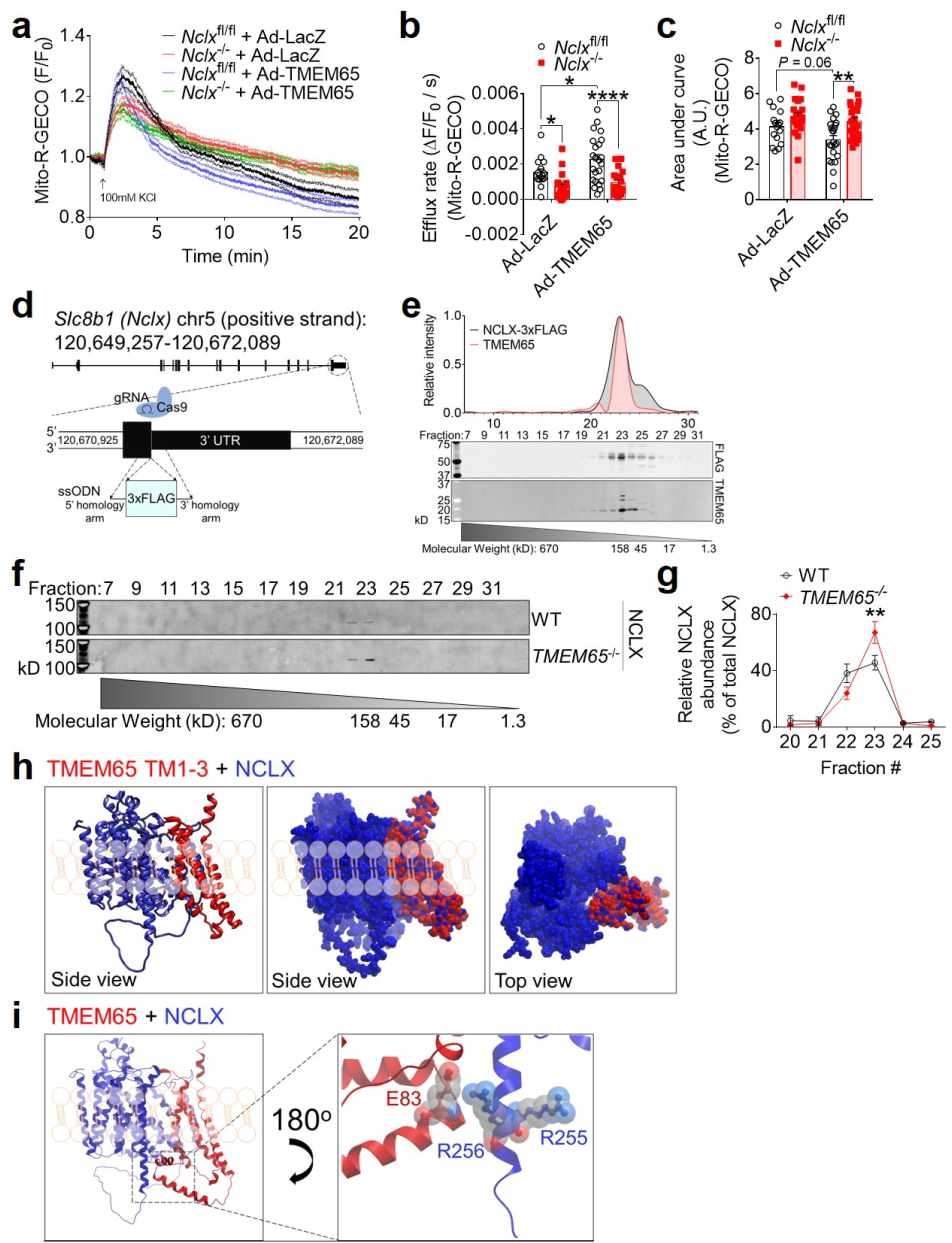
TMEM65 and NCLX interact functionally and physically. a, Pooled traces showing KCl-evoked _m_Ca^2+^ transients as measured by Mito-R-GECO fluorescence in intact *Nclx*^fl/fl^ and *Nclx*^-/-^ mouse embryonic fibroblasts after transduction with Ad-LacZ or Ad-TMEM65 (*n* = 16 cells for *Nclx*^fl/fl^ + Ad-LacZ; 21 for *Nclx*^-/-^ + Ad-LacZ; 23 for *Nclx*^fl/fl^ + Ad-TMEM65; 24 for *Nclx*^-/-^ + Ad-TMEM65). _m_Ca^2+^ efflux rate measured over the first 60 seconds following the peak of the mitochondrial Ca^2+^ transient (b), and area under the curve (c) for _m_Ca^2+^ transients shown in panel (a). (*n* = 16 cells for *Nclx*^fl/fl^ + Ad-LacZ; 21 for *Nclx*^-/-^ + Ad-LacZ; 23 for *Nclx*^fl/fl^ + Ad-TMEM65; 24 for *Nclx*^-/-^ + Ad-TMEM65; \**P*<0.05; \*\**P*<0.01; \*\*\*\**P*<0.0001 by 2-way ANOVA with Sidak’s post hoc test). d, CRISPR-Cas9 knock-in strategy to insert a 3xFLAG epitope tag after the final sense codon of the mouse *Slc8b1* gene. ssODN, single-stranded oligodeoxynucleotide. e, Western blotting showing molecular weight fraction distribution of NCLX-3xFLAG and TMEM65 after size-exclusion chromatography of isolated mouse heart mitochondria. Chromatogram and molecular weight standard curve for gel filtration standards of known molecular weight are shown in Extended Data Fig. 3d-e. f, Western blotting showing molecular weight fraction distribution of endogenous NCLX after size-exclusion chromatography of isolated WT and *TMEM65*^-/-^ AC16 cardiomyocyte mitochondria. g, quantification of NCLX signal intensity across fractions shown in panel (f), normalized to total NCLX signal intensity. (*n* = 3 / genotype; \*\**P*<0.01 by 2-way ANOVA with Sidak’s post-hoc test. h, *In silico* molecular modeling showing predicted interaction between NCLX (blue) and the transmembrane domains (TM1-3) of TMEM65 (red). Modeling places NCLX adjacent to the longest transmembrane helix of NCLX, depicted in ribbon and space-filling views. i, Superimposing the structure of full-length, mature mitochondrial TMEM65 onto the model shown in panel (h) predicts electrostatic interaction between the soluble domain of TMEM65 and the positively-charged regulatory region of NCLX’s longest transmembrane helix that includes R255 and R256.

**Extended Data Fig. 3:**
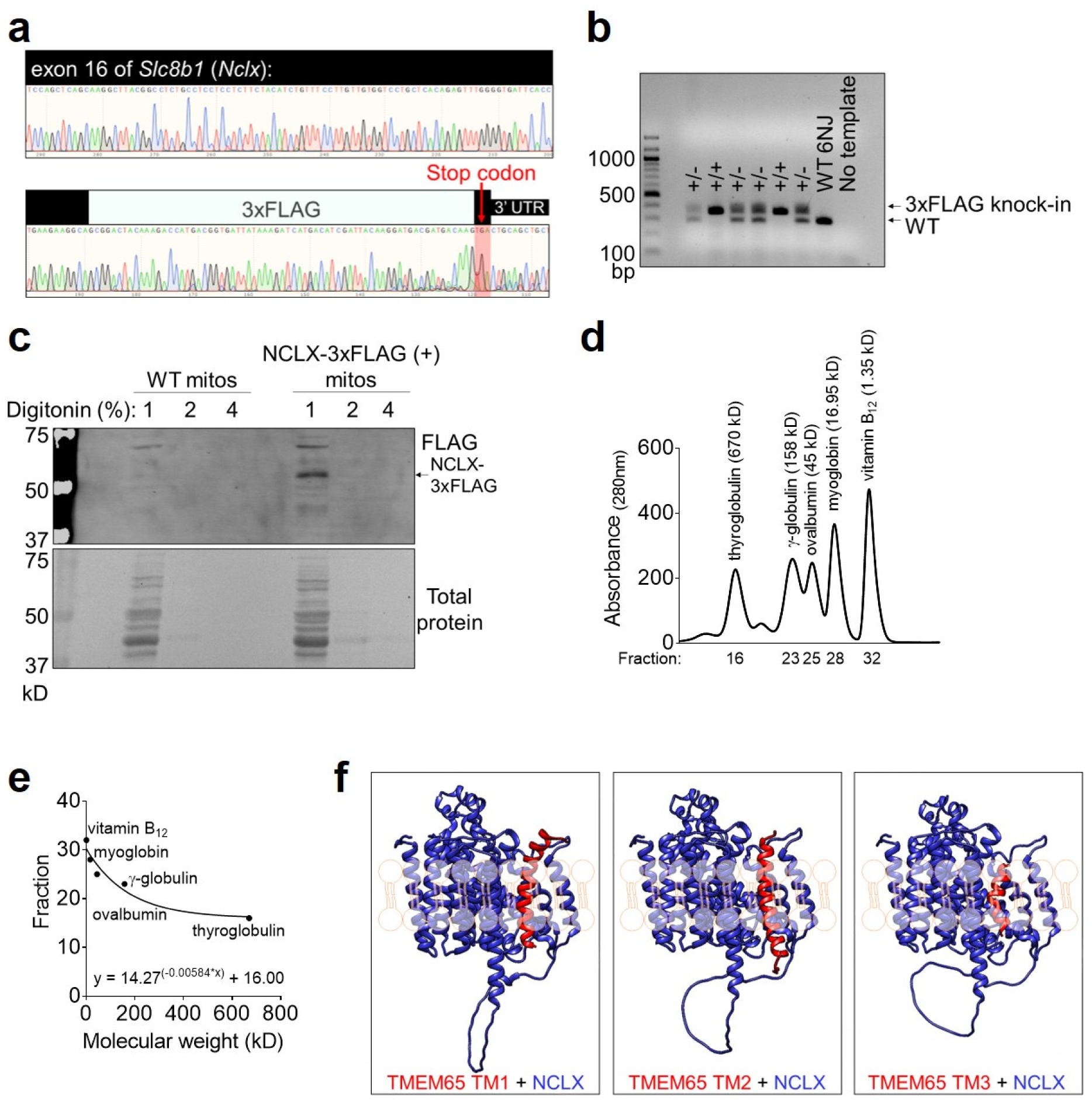
Validation of NCLX-3xFLAG mouse and stages in modeling TMEM65-NCLX interaction. a, Chromatogram of sequencing confirming proper in-frame insertion of DNA encoding the NCX-3xFLAG epitope tag prior to the stop codon of the endogenous mouse *Nclx* gene. b, Genotyping gel showing pups heterozygous (+/-) or homozygous (+/+) for the 3xFLAG knock-in. c, Western blots validating solubilization and specific detection of NCLX-3xFLAG protein (arrow) in isolated adult mouse heart mitochondria. Amido black staining for total protein is shown as a loading control. d, Absorbance at 280nm for gel filtration standards separated by size-exclusion chromatography. e, Standard curve of elution fraction vs. known molecular weight for gel filtration standards separated by size-exclusion chromatography. The equation shows the fit of the standards to a one-phase exponential decay. f, *In silico* molecular modeling showing interactions between individual TMEM65 transmembrane (TM) domains (red) and NCLX (blue).

## NCLX and TMEM65 interactions and structural prediction of a macromolecular complex

To facilitate *in situ* investigative approaches for NCLX and its interacting partners, we generated a NCLX-3xFLAG knock-in mouse model, in which a C-terminal 3xFLAG epitope tag was knocked into exon 16 of *Slc8b1* (Fig. 3d, Extended Data Fig. 3a-c). Fast protein size-exclusion liquid chromatography (FPLC) of mitochondria isolated in non-reducing conditions from the hearts of NCLX-3xFLAG knock-in mice showed that the distribution of protein complexes containing NCLX mirrored that of TMEM65, with the abundance of both proteins greatest in the ∼160 kD containing fraction (Fig. 3e; Extended Data Fig 3d-e). The distribution pattern was consistent with NCLX and TMEM65 existing in a common macromolecular complex. To directly test this notion and circumvent the technical challenges associated with immunoprecipitating NCLX, we performed size-exclusion FPLC on non-reduced mitochondrial samples isolated from WT and *TMEM65*^-/-^ AC16 cardiomyocytes (Fig. 3f). Under these experimental conditions, NCLX from WT mitochondria fractionated as a stable dimer >100kD, which was evenly distributed between elution fractions 22 and 23 (∼140-kD) (Fig. 3f; Extended Data Fig. 3e). This reaffirms previous reports of NCLX, and other sodium/calcium exchangers, forming SDS resistant dimers^1^. The molecular weight of mature mitochondrial human NCLX is ∼61-kD, and the molecular weight of mature mitochondrial human TMEM65 is ∼19-kD. Thus, this elution pattern of NCLX at a molecular mass ∼140-kD aligns closely with a predicted macromolecular complex consisting of 2 NCLX: 1 TMEM65. Fitting with this model, genetic deletion of *TMEM65* resulted in a right-ward shift in the NCLX distribution, with a significant increase in relative NCLX abundance in fraction 23 (∼120-kD, Fig. 3f-g), corresponding to the weight of an NCLX homodimer lacking TMEM65.

We then turned to *in silico* modeling using the predicted AlphaFold structures of human TMEM65 and NCLX to interrogate how these proteins might physically interact. Modeling of interactions between each of the three individual TMEM65 transmembrane domains with NCLX yielded a consensus model, in which each individual transmembrane domain of TMEM65 aligned within the same cleft next to the longest transmembrane helix of NCLX, which feeds into its soluble loop domain (Extended Data Fig. 3f). Intriguingly, modeling of interactions between NCLX and a truncated TMEM65 consisting of all 3 of its transmembrane domains likewise placed TMEM65 in this same cleft on NCLX (Fig. 3h). Superposition of full length, mature mitochondrial TMEM65 onto this alignment places the soluble N-terminal ‘matrix hook domain’ of TMEM65 in a position to interact with both the soluble ‘regulatory loop’ of NCLX and the C-terminal end of NCLX’s longest transmembrane helix (Fig 3i). The C-terminal end of this NCLX region contains a cluster of positively charged arginine residues (R253, R255, R256) that are reported to limit NCLX activity as mitochondria depolarize^22^. We hypothesize that interactions between negatively-charged residues in this soluble hook domain of TMEM65, such as glutamic acid 83 (Fig 3i), and this positively charged regulatory region of NCLX ultimately enhance NCLX activity, as has been reported to occur with the introduction of a local negative charge by inserting a phosphomimic mutation in the nearby NCLX residue, S258^23^.

## Knockdown of *Tmem65* promotes pathogenic _m_Ca^2+^ overload and striated muscle pathology *in vivo*

Disruption of NCLX-dependent _m_Ca^2+^ efflux predisposes to catastrophic _m_Ca^2+^ overload, cell death, and progressive organ-level dysfunction in excitable tissues such as the heart and the brain^3,4^. Based on our conclusion that TMEM65 promotes NCLX activity, we reasoned that genetic disruption of *Tmem65* expression should recapitulate these aspects of NCLX disruption *in vivo*. Indeed, adeno-associated virus 9 (AAV9)-delivery of shRNA targeting *Tmem65* in mice reduced TMEM65 expression and decreased inhibitory PDH-S293 phosphorylation in skeletal muscle (Fig. 4a-c), indicating that reduced TMEM65 expression is sufficient to perturb _m_Ca^2+^ efflux and increase _m_Ca^2+^ loading in intact tissue. We noted that *Tmem65* knockdown (KD) produced a robust neuromuscular phenotype evident as early as 3 weeks of age. *Tmem65* KD mice exhibited an altered, waddling gait and abnormal posturing of the hindlimbs when suspended by the tail (Fig. 4d), classical signs of neuromuscular dysfunction. In line with this phenotype, *Tmem65* KD mice displayed a diminished functional capacity to hold on to an inverted wire grid when assessed via 4-limb wire hang test (Fig. 4e). Skeletal muscle fiber size and muscle mass (Fig 4f-g; Extended Data Fig. 4a-b) were reduced in *Tmem65* KD animals. Furthermore, the percentage of centrally-nucleated skeletal muscle fibers, an index of recent fiber death and regeneration, tended to increase with loss of TMEM65 (Fig. 4h). We likewise noted reduced TMEM65 expression and decreased PDH phosphorylation in the hearts of *Tmem65* KD mice (Fig. 4i-k), and diminished cardiac function as early as 6 weeks of age (Fig. 4l; Extended Data Fig. 4c-d), which correlated pairwise with the loss of TMEM65 protein expression (Fig. 4m). These observations suggest that reduced TMEM65 expression elicits pathogenic _m_Ca^2+^ overload *in vivo* that increases cell death and striated muscle dysfunction.

**Figure 4:**
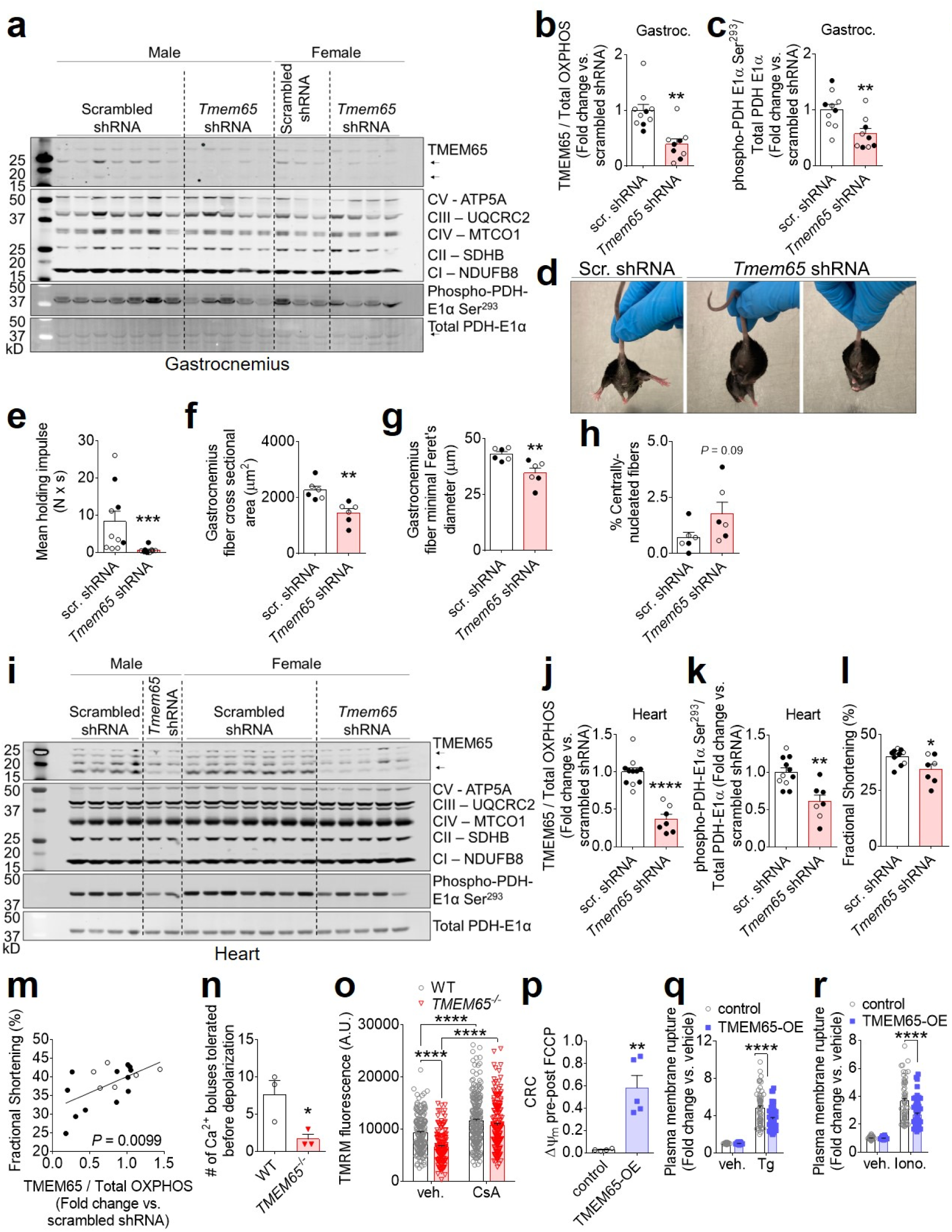
Loss of TMEM65 expression impairs striated muscl function and predisposes to _m_Ca^2+^-overload-induced cell death, while TMEM65 overexpression protects against Ca^2+^-induced cell death. a, Western blots confirming reduction in TMEM65 protein expression in gastrocnemius muscle of 24-week-old mice injected as neonates with AAV9-*Tmem65* shRNA. Dashed lines are to aid in visualizing separate conditions. b, Densitometric quantification of gastrocnemius muscle TMEM65 expression, normalized to total OXPHOS complexes (CI-CV). Male mice are shown in open symbols, female mice are shown in filled symbols. scr., scrambled. (*n* = 10 mice + scrambled shRNA, 9 mice + *Tmem65* shRNA, \*\**P*<0.01 by Mann-Whitney test). c, Densitometric quantification of gastrocnemius PDH phosphorylation normalized to total PDH. (*n* = 10 mice + scrambled shRNA, 9 mice + *Tmem65* shRNA, \*\**P*<0.01 by unpaired t-test). d, Representative images showing abnormal hindlimb positioning and/or clasping in *Tmem65* knockdown (KD) mice at 14 weeks of age. scr., scrambled. e, Mean holding impulse representing performance of mice on 4-limb wire hang test at 24 weeks of age. (*n* = 10 mice + scrambled shRNA, 9 mice + *Tmem65* shRNA. \*\*\**P*<0.001 by Mann-Whitney test). Fiber cross-sectional-area (f), minimal Feret’s diameter (g), and % centrally nucleated fibers (h) for the gastrocnemius muscle. (*n* = 6 mice / group. \*\**P*<0.01 by unpaired t-test). i, Western blots confirming reduction in TMEM65 protein expression in the hearts of *Tmem65* KD mice at 6 weeks of age. Dashed lines are to aid in visualizing separate conditions. Densitometric quantification of heart TMEM65 expression, normalized to total OXPHOS complexes (CI-CV) (j), and densitometric quantification of heart PDH phosphorylation normalized to total PDH (k) (*n* = 11 mice + scrambled shRNA, 7 mice + *Tmem65* shRNA. \*\**P*<0.01, \*\*\*\**P*<0.0001 by unpaired t-test). l, Left ventricular fractional shortening measured by echocardiography in mice at 6 weeks of age (*n* = 11 mice + scrambled shRNA, 7 mice + *Tmem65* shRNA. \**P*<0.05 by unpaired t-test). m, Linear regression showing positive correlation between left ventricular fractional shortening and TMEM65 protein expression, as measured in mice with scrambled shRNA (*n* = 11) and mice with *Tmem65* shRNA (*n* = 7). n, Number of Ca^2+^ boluses tolerated prior to spontaneous mitochondrial depolarization and release of matrix Ca^2+^ for the calcium retention capacity assay in permeabilized AC16 cells shown in Extended Data Fig. 4e (*n* = 3/genotype. \**P*<0.05 by unpaired t-test). o, TMRM staining intensity as a marker of mitochondrial membrane potential in cells incubated acutely with vehicle (veh.) or cyclosporine A (CsA). (*n* = 170 cells for WT + vehicle; 140 cells for *TMEM65*^-/-^ + vehicle; 225 cells for WT + CsA; 178 cells for TMEM65^-/-^ + CsA. \*\*\*\**P*<0.0001 by two-way ANOVA with Sidak’s post-hoc test). p, Magnitude of drop in JC-1 fluorescence ratio upon addition of FCCP, corresponding to mitochondrial membrane potential after the final Ca^2+^ addition, for calcium retention capacity (CRC) assay in permeabilized AC16 cells shown in Extended Data Fig. 4f. (*n* = 4 replicates for empty vector control, 5 replicates for TMEM65-OE; \*\**P*<0.01 by unpaired t-test with Welch’s correction). q, Plasma membrane rupture as measured by Sytox green staining, as an indicator of cell death in response to 24-hour treatment with thapsigargin (Tg). (*n* = 56 for control + vehicle; 60/group for control + Tg, TMEM65-OE + vehicle, and TMEM65-OE + Tg; \*\*\*\**P*<0.001 by 2-way ANOVA with Sidak’s post-hoc test). r, Plasma membrane rupture in response to 24- hour treatment with ionomycin (Iono.) (*n* =55 for control + vehicle; 60/group for control + Iono., TMEM65-OE + vehicle, and TMEM65-OE + Iono.; \*\*\*\**P*<0.001 by 2-way ANOVA with Sidak’s post-hoc test).

**Extended Data Fig. 4:**
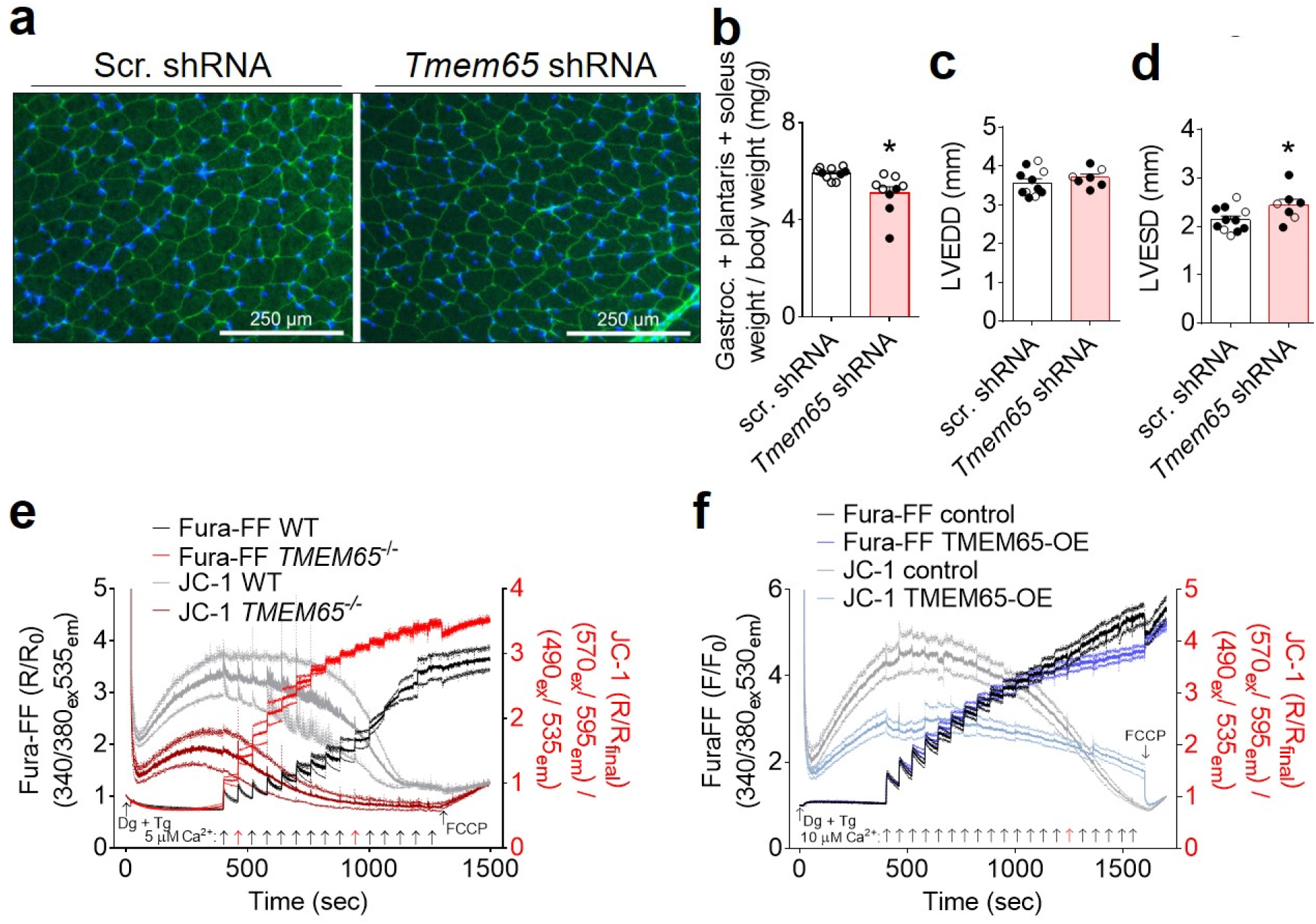
Phenotypic consequences of altered TMEM65 expression. a, Representative images showing WGA labelling (green) and nuclear staining with DAPI (blue) in cross sections of the gastrocnemius muscle at 24 weeks of age. scr., scrambled. b, Total mass of the gastrocnemius + soleus + plantaris muscle, dissected as a unit, normalized to total body mass at 24 weeks of age. Open symbols represent male mice, filled symbols represent female mice. (*n* = 10 mice + scrambled shRNA, 9 mice + *Tmem65* shRNA, \**P*<0.05 by unpaired t-test with Welch’s correction). Left ventricular end diastolic dimension (LVEDD) (c) and left ventricular end systolic dimension (LVESD) (d) as measured by echocardiography in mice at 6 weeks of age (*n* = 11 mice + scrambled shRNA, 7 mice + *Tmem65* shRNA. \**P*<0.05 unpaired t-test). e, Pooled traces showing extramitochondrial bath Ca^2+^ (Fura-FF) and mitochondrial membrane potential (JC-1) in permeabilized WT and *TMEM65*^-/-^ AC16 cardiomyocytes in response to repeated additions of 5µM Ca^2+^ (arrows). Red arrows indicate the Ca^2+^ bolus typically triggering permeability transition in each genotype. (*n* = 3 replicates/genotype). f, Pooled traces showing extramitochondrial bath Ca^2+^ (Fura-FF) and mitochondrial membrane potential (JC-1) in permeabilized control or TMEM65-OE AC16 cardiomyocytes in response to repeated additions of 10µM Ca^2+^ (arrows). Red arrow indicates Ca^2+^ bolus triggering permeability transition in control cells. (*n* = 4 replicates for empty vector controls, 5 replicates for TMEM65-OE).

## Loss of TMEM65 predisposes to cell death and gain of TMEM65 protects against Ca^2+^- induced mitochondrial permeability transition and cell death

To mechanistically investigate the relationship between TMEM65 expression, _m_Ca^2+^ overload, and cell death, we next challenged permeabilized AC16 cells with repeated boluses of Ca^2+^. Wild-type cells exhibited a precipitous drop in mitochondrial membrane potential (ΔΨ_m_), accompanied by spontaneous release of _m_Ca^2+^, after the delivery of ∼ 10 Ca^2+^ boluses, whereas *TMEM65*^-/-^ cells depolarized and released _m_Ca^2+^ with fewer Ca^2+^ boluses (Fig. 4n, Extended Data Fig. 4e). TMRM staining of intact cells revealed that loss of TMEM65 slightly reduced resting ΔΨ_m_, and that this effect could be rescued by acute incubation with the mitochondrial permeability transition pore (mPTP) inhibitor cyclosporin A (CsA) (Fig. 4o). These findings suggest that similar to genetic loss of NCLX, loss of TMEM65 increases the susceptibility to ongoing _m_Ca^2+^ overload, leading to increased mPTP activity that can compromise mitochondrial function and culminate in cell death.

Intriguingly, in calcium retention capacity assays TMEM65-OE modestly reduced net _m_Ca^2+^ uptake with each successive Ca^2+^ bolus and resulted in a gradual decline in ΔΨ_m_. However, TMEM65-OE completely prevented the full depolarization of ΔΨ_m_ and the abrupt release of _m_Ca^2+^ characteristic of mitochondrial permeability transition with the administration of up to 20 Ca^2+^ boluses (Fig. 4p, Extended Data Fig. 4f). This result bears a striking resemblance to that expected with enhanced, electrogenic (3 Na^+^ in : 1 Ca^2+^ out) forward-mode NCLX activity. Finally, we assessed the pathophysiological relevance of increased TMEM65 expression by assessing its effects on thapsigargin-or ionomycin-induced cell death in AC16 cardiomyocytes as a model for cytosolic Ca^2+^ and subsequent _m_Ca^2+^ overload. TMEM65-OE was sufficient to limit cell death in response to 24-hour thapsigargin or ionomycin treatment (Fig. 4q-r), indicating the power of enhanced TMEM65 expression to limit pathogenic _m_Ca^2+^ stress.

The maintenance of _m_Ca^2+^ homeostasis represents a critical challenge, as cells in energetically demanding tissues depend on _m_Ca^2+^-dependent regulation of metabolism to match ATP production to fluctuations in cellular activity and ATP consumption. At the same time, these cells must avoid excessive _m_Ca^2+^ accumulation or risk mitochondrial permeability transition and cell death, a key feature observed in both acute and chronic pathologies such as ischemia-reperfusion injury, chronic heart failure, and muscular dystrophies. Prior *in vivo* studies have revealed the efficacy of enhancing NCLX expression and net _m_Ca^2+^ efflux in limiting the progression of heart failure following myocardial infarction or sustained pressure-or neurohormonal overload^3,5^, and in slowing the progression of cognitive decline and neurodegeneration in experimental models of Alzheimer’s disease^6^. Therapeutically enhancing NCLX activity has emerged as a desirable goal for the treatment of such diseases. Indeed, this approach may offer benefits over strategies designed to attenuate _m_Ca^2+^ uptake through the mitochondrial calcium uniporter channel, as it may prevent the progressive accumulation of deleterious levels of _m_Ca^2+^ over time without compromising acute and transient elevations in _m_Ca^2+^ that may contribute to the physiologic regulation of mitochondrial energetics. However, no chemical compounds nor endogenous regulators that can specifically enhance NCLX activity have yet been described.

Our *in vitro* findings identify TMEM65 as a potent enhancer of _m_Ca^2+^ efflux and indicate that it mediates this function by positively regulating its interacting partner, NCLX. The observations that TMEM65 overexpression increases and that *TMEM65* knockout diminishes Na^+^-sensitive _m_Ca^2+^ efflux support this conclusion, as NCLX is the only known mitochondrial Na^+^/Ca^2+^ exchanger. Moreover, we find that genetic deletion of NCLX mitigates TMEM65-dependent _m_Ca^2+^ efflux, indicating that TMEM65 enhances _m_Ca^2+^ efflux through, rather than independent of, NCLX. Our identification of TMEM65 within the NCLX interactome and our finding that the molecular weight of macromolecular protein complexes containing NCLX is reduced in the absence of TMEM65 support a model in which TMEM65 physically interacts with NCLX to regulate its function. TMEM65 has no known enzymatic activity nor domains resembling kinases or other regulatory enzymes; therefore, we hypothesize that TMEM65 exerts its effects on NCLX-dependent _m_Ca^2+^ efflux through allosteric regulation of NCLX. Our *in silico* modeling of a TMEM65-NCLX complex proposes inter-molecular interactions between the soluble domain of TMEM65 and a known regulatory helix of NCLX^22^ that may mediate enhancement of NCLX activity.

The neuromuscular and cardiac dysfunction that develops with only partial knockdown of *Tmem65* expression *in vivo* demonstrates the importance of this protein within excitable tissues. The additional observation that reducing TMEM65 expression in skeletal muscle and heart tissue results in pathogenic _m_Ca^2+^ overload reinforces the conclusion that TMEM65 has a fundamental role in maintaining physiologic _m_Ca^2+^ homeostasis in the intact organism. It is notable that homozygous null mutation of *TMEM65* in humans causes a mitochondrial encephalomyopathy with neurological deficits, muscle weakness, and impaired mitochondrial energetics^17^, all phenotypes that are consistent with pathologic _m_Ca^2+^ overload. Complete genetic loss of NCLX is embryonic lethal^3^, precluding study of its functional consequences *in vivo*. However, we propose that TMEM65 is an essential positive regulator of NCLX and that the human *TMEM65*^-/-^ phenotype described by Nazli et al^17^ and the effects of *Tmem65* knockdown we report here result from impairment of NCLX function. Finally, our finding that TMEM65 overexpression *in vitro* is sufficient to limit _m_Ca^2+^ overload-induced cell death highlights TMEM65 as a novel translational target for diseases driven by pathogenic _m_Ca^2+^ accumulation and cell dropout.

Our discovery of TMEM65 as the first genetically-confirmed regulator of NCLX provides mechanistic insight into the endogenous modulation of _m_Ca^2+^ efflux and opens an avenue for the design of strategies to modulate NCLX activity that have thus far eluded the field. We further demonstrate the physiological relevance of TMEM65 in mitigating deleterious _m_Ca^2+^ overload *in vivo* and *in vitro*, and conclude that modulation of TMEM65 and/or its interaction with NCLX could be an innovative approach for the therapeutic control of mitochondrial calcium homeostasis in human disease.

## Methods

### Generation of NCLX-BioID2-HA fusion construct

Human *SLC8B1* cDNA (Origene #RC214624; NM_024959) was PCR amplified using primers to introduce a 5′ AgeI restriction site and a 3′ BamHI restriction site flanking the coding sequence. The PCR product was cloned via AgeI/BamHI into an MCS-BioID2-HA plasmid (Addgene #74224)^15^ to enable expression of an NCLX-BioID2-HA fusion protein under the control of a CMV promoter. The sequence encoding BioID2-HA was PCR amplified out of the MCS-BioID2- HA plasmid, using a forward primer to introduce a start codon immediately after the 5’ BamHI restriction site, and a reverse primer including the 3’ HinDIII site. The PCR product was cloned back into the MCS-BioID2-HA backbone via BamHI/HinDIII to enable expression of the BioID2- HA protein alone as a negative control for non-specific protein biotinylation.

### Cell culture and transfection

AC16 cardiomyocytes (EMD Millipore #SCC109) were maintained in DMEM/F12 (Thermo Scientific #11320082) supplemented with 12.5% fetal bovine serum (Peak Serum #PS-FB3) + 1% penicillin/streptomycin (Sigma-Aldrich #P0781-100ML). Cells were grown to 75% confluency, then washed in 1x PBS and switched to Opti-MEM reduced serum media (Thermo Fisher Scientific #31985070). After 1 hour, plasmids were transiently transfected into cells using Lipofectamine 3000 (Invitrogen #L3000015) in Opti-MEM according to the manufacturer’s instructions, and allowed to express for 24 hours before analysis.

### Cellular fractionation and mitochondrial isolation

Cells were washed in 1x phosphate buffered saline (PBS) (Morganville Scientific #PH0200), scraped off of culture dishes into fresh PBS, then centrifuged for 5 min at 200g at 4°C. Pelleted cells were resuspended in isotonic mitochondrial isolation buffer (10mM HEPES, 200mM mannitol, 70mM sucrose, 1mM ethylene glycol-bis(β-aminoethyl ether)-N,N,N′,N′-tetraacetic acid (EGTA), pH = 7.5) and homogenized in a handheld glass dounce homogenizer. Homogenates were centrifuged at 500g for 10min at 4°C to remove insoluble material. An aliquot of supernatant was saved as whole cell lysate, and the remaining supernatant was centrifuged at 12,000g for 15min at 4°C to pellet mitochondria. The supernatant was transferred to a new tube and centrifuged again at 12,000g for 15min at 4°C to remove any residual mitochondria, and then saved as the cytosolic fraction. Pelleted mitochondria were washed twice in fresh isotonic mitochondrial isolation buffer followed by centrifugation at 12,000g for 15min at 4°C. The final mitochondrial pellet was resuspended as indicated for use in the assays described below.

### Sub-mitochondrial protein localization

Mitochondria isolated from AC16 cardiomyocytes transiently transfected with NCLX-BioID2-HA were resuspended in intracellular buffer (120 mM KCl, 10 mM NaCl, 1 mM KH_2_PO_4_, 20 mM HEPES-Tris, pH 7.2). Trypsin was added at concentrations from 0-0.25% and mitochondria were incubated at 37°C for 7.5min. Samples were placed on ice, and the digestion was stopped by adding SIGMAFAST protease inhibitor cocktail (Sigma-Aldrich #S8830-20TAB) and 5x SDS sample buffer (250µM Tris-HCl, pH 7.0; 40% glycerol; 8% sodium dodecyl sulfate; 20% β-mercaptoethanol; 0.1% bromophenol blue) to a 1x final concentration and boiling the samples at 95°C for 5min prior to analysis via western blotting.

### Proximity biotinylation assay

Biotin (Sigma-Aldrich #B4501-1G) was added to the culture media at a final concentration of 50µM at the time of transfection. After 24 hours, cells were washed in 1x PBS and trypsinized from cell culture plates in 0.25% trypsin EDTA (Corning #25-053-CI), centrifuged for 5min at 200g, and washed twice in 1x PBS. Cell pellets were snap frozen in liquid nitrogen until analysis. One-tenth of each sample was reserved for western blotting to confirm expression of BioID2-HA constructs and lysed in BioID lysis buffer (8M urea, 50mM Tris-Cl pH 7.4, 1mM DTT, 3.6% Triton-X-100 + 1x SIGMAFAST protease inhibitor cocktail), sonicated for 10sec, rotated for 30min at 4°C, then centrifuged at 5000g for 5 min at 4°C to remove insoluble material. The remainder of each sample was submitted to the Sanford Burnham Prebys Proteomics Core for affinity purification of biotinylated proteins followed by identification via mass spectrometry, as described previously^15^. Biotinylated proteins enriched at least 2-fold in NCLX-BioID2-HA samples versus BioID2-HA samples in at least 2/3 replicates, with known or predicted mitochondrial localization according to the 2018 release of the Integrated Mitochondrial Protein Index database (https://www.mrc-mbu.cam.ac.uk/research-resources-and-facilities/impi), were considered as NCLX proximal proteins.

### Western blotting

Protein concentration in sample lysates was determined by bicinchoninic acid protein assay (bioWORLD #20831001-1). 5x SDS sample buffer was added to a 1x final concentration. 20-50 µg of protein lysates were loaded to 10% or 12% polyacrylamide Tris-glycine SDS gels or to large-format NuPAGE 4-12% gradient Bis-Tris gels (Thermo Fisher Scientific # WG1403BOX and # WG1402BOX). After separation by electrophoresis, proteins were transferred to polyvinyldienefluoride membranes (Millipore #IPFL00010) and subjected to western blotting as described previously^5^.

Primary antibodies and their dilutions were: rat monoclonal against HA (Roche #11867423001), 1:1000; mouse monoclonal against VDAC1 (Abcam #ab14734), 1:1000; rabbit monoclonal against MCU (Cell Signaling Technology #14997), 1:1000; mouse monoclonal anti-FLAG (Sigma-Aldrich #F1804-1MG), 1:200; rabbit polyclonal against TMEM65 (Abcam #ab236861), 1:1000); rabbit monoclonal against Tom20 (Cell Signaling Technology #42406S), 1:1000; mouse monoclonal against cyclophilin D (Abcam #110324), 1:1000; rabbit polyclonal against Myc (Invitrogen #PA1-981), 1:1000; mouse monoclonal against tubulin (Abcam #ab7291), 1:1000; mouse monoclonal total OXPHOS rodent antibody cocktail (Abcam #ab110413), 1:1000; rabbit polyclonal against NCLX (Aviva Systems Biology #ARP44042_P050), 1:500; rabbit polyclonal against NCLX (Abcam # ab83551), 1:500; rabbit polyclonal against phospho-PDH E1α-Ser293 (Abcam #ab92696) 1:1000; and mouse monoclonal against PDH E1α subunit (Abcam #ab110330) 1:1000. IRDye 800CW streptavidin (LI-COR #925-32230) was used at a dilution of 1:5000. Secondary antibodies and their dilutions included: IRDye 680RD goat anti-rabbit (LI-COR #926-68071), 1:10,000; IRDye 800CW goat anti-mouse (LI-COR #926-32210), 1:10,000; IRDye 800CW goat anti-rabbit (LI-COR #925-32210), 1,10,000; IRDye 680RD goat anti-mouse (LI-COR #925-68070), 1:10,000; and IRDye 680RD Goat anti-Rat (LI-COR # 925- 68076), 1:10,000.

For visualization of total protein loading, membranes were stained with amido black (Fisher Scientific #AAA1137414) as described previously^24^ and imaged on a Gel Dox XR+ Molecular Imager (Bio-Rad). Western blot densitometry was measured using LI-COR Image Studio software (LI-COR, version 2.0.38). All full-length western blots are shown in Supplemental Figures.

### Confirmation of TMEM65 mitochondrial localization

AC16 cardiomyocytes grown on 35mm glass-bottomed plates (MatTek #P35G1.510C) were transiently transfected as described above with a plasmid encoding C-terminal Myc-FLAG epitope-tagged human TMEM65 (TMEM65-Myc-FLAG) (Origene #RC207368; NM_194291). Cells were transfected with empty pCMV6-Entry vector (Origene #PS100001) as a negative control. After 24 hours, the cells were fixed in 3% paraformaldehyde (Fisher Scientific #T353- 500) in 1xPBS for 15 min at room temperature, then washed 3 times for 5min in 1xPBS. Fixed cells were blocked and permeabilized overnight at 4°C in block solution (1x PBS + 5% bovine serum albumin (Sigma-Aldrich #A3803) + 0.5% Triton-X-100). Cells were then incubated in primary antibodies diluted in block solution for 1.5 hours at room temperature, washed 3 times in 1xPBS, and incubated in secondary antibodies diluted in block solution for 1 hour at room temperature in the dark. Cells were washed 3 times in 1x PBS and mounted with glass coverslips using ProLong Gold Antifade Mountant with DAPI (Invitrogen #P36935) before imaging on a Zeiss LSM 900 microscope with an Airyscan 2 detector. Primary antibodies and dilutions included mouse monoclonal against FLAG at 1:200, and rabbit polyclonal against Tom20 at 1:200 as a mitochondrial marker. Secondary antibodies and dilutions included goat anti-mouse IgG-AlexaFluor 594 (Invitrogen #A-11005) at 1:1000 and goat anti-rabbit IgG-AlexaFluor 488 (Invitrogen #A-11008) at 1:1000.

In separate experiments, AC16 cardiomyocytes were transfected with the plasmid encoding TMEM5-Myc-FLAG or empty vector and used for cellular fractionation and mitochondrial isolation. Mitochondria were resuspended in isotonic mitochondrial isolation buffer, and the whole cell lysate, cytosolic fraction, and mitochondrial fraction were subjected to western blotting.

### qPCR analysis of mouse *Tmem65* gene expression

Tissues were collected from adult male and female C57BL/6NJ mice (Jackson Laboratory #005304) at 10-15 weeks of age. RNA was isolated and cDNA prepared, then subjected to qPCR as described elsewhere^5^. qPCR primers against mouse transcripts were: *Tmem65* forward: 5’- GCGACTTCATCTACAGCCTGCA-3’ and *Tmem65* reverse: 5’- TGTCCTGGAGTGGGTGGTAGAG-3’; and *Rps13* forward: 5’- GCACCTTGAGAGGAACAGAA- 3’ and *Rps13* reverse: 5’- GAGCACCCGCTTAGTCTTATAG-3’.

### Generation of cell lines with stable TMEM65 overexpression

AC16 cardiomyocytes were transfected with the TMEM65-Myc-FLAG plasmid or empty vector. After 48 hours, cells were returned to complete medium and selected with G418 (ThermoScientific # 10131035) at 600µg/mL until no cells survived on non-transfected control plates. Surviving cells were clonally isolated using glass cloning cylinders (Corning # 3166-6). To confirm stable expression of TMEM65-Myc-FLAG, cells were lysed in 1x RIPA buffer (EMD Millipore #20-188) supplemented with 1x SIGMAFAST protease inhibitor cocktail and 1X Phosstop phosphatase inhibitor (Roche #04906837001), sonicated for 10sec, and incubated on ice for 30min. Lysates were centrifuged at 5000g for 5 min at 4°C, and the supernatant was used for western blotting. TMEM65-Myc-FLAG stable overexpression line #20 (“TMEM65-OE”) was used throughout.

### Generation of stable *TMEM65* knockout cells

AC16 cardiomyocytes were changed to Opti-MEM and transfected with a custom plasmid (VectorBuilder) encoding human codon-optimized SpCas9 under the control of a CBh promoter, two gRNAs under the control of U6 promoters targeting exon 1 of human *TMEM65* (5’- GCCGTGTTCAGCGCCTCCAT-3’ (chr8:124371952-124371971); and 5’-CGCGCCCCCCGGCGGCTTGC-3’ (chr8:124372016-124372035)), and a blasticidin resistance gene under the control of a hPGK promoter, using FuGENE HD Transfection Reagent (Promega #E2311) according to the manufacturer’s instructions. Non-transfected wild-type (WT) controls were switched to Opti-MEM in parallel. After 48hrs, cells were returned to complete media, and transfected cells were selected with blasticidin (Invivogen #ant-bl-05) at 10µg/mL until no cells survived on non-transfected control plates. Surviving stably-transfected cells were clonally isolated using glass cloning cylinders. Loss of TMEM65 protein in clonal lines was confirmed via western blot. *TMEM65*^-/-^ lines #2 and #5 were used throughout.

### Generation of immortalized *Nclx*^fl/fl^ mouse embryonic fibroblasts (MEFs) and acute deletion of *Nclx*

Fibroblasts were isolated from E13.5 *Nclx*^fl/fl^ mouse embryos^3^ and cultured as described previously^25^. MEFs were immortalized by addition of SV40 T antigen lentivirus (Alstem #C1LV01) in the presence of 0.02 mg/mL polybrene (EMD Millipore # TR-1003-G), according to the manufacturer’s instructions. Immortalized MEFs were transduced for 24 hours with Ad-LacZ/Ad-β-gal adenovirus (Vector Biolabs #1080) under the control of a CMV promoter as a control, or adenovirus encoding Cre recombinase (Ad-Cre) (Vector Biolabs # 1045) under the control of a CMV promoter to delete NCLX (“*Nclx*^-/-^”). Experiments were performed 7 days later to allow sufficient time for protein turnover.

### Adenoviral gene transfer

AC16 cardiomyocytes and MEFs were cultured as described above. C2C12 skeletal myoblasts (ATCC #CRL1772) were grown in DMEM (Corning # 15-013-CV) supplemented with 10% fetal bovine serum and 1% penicillin/streptomycin. Cells were transduced as indicated for 24-48 hours with custom adenovirus encoding untagged human TMEM65 (Ad-TMEM65) under the control of a CMV promoter at 100-500 multiplicity of infection (MOI). For negative controls, cells were transduced with adenovirus encoding eGFP (Ad-eGFP) (VectorBuilder # VB010000- 9299hac) or LacZ/β-galactosidase (Ad-LacZ/Ad-β-gal) under the control of a CMV promoter at an equivalent MOI.

### Mitochondrial Ca^2+^ flux assays

Cells were trypsinized from culture plates and used for assessment of _m_Ca^2+^ efflux and mitochondrial Ca^2+^ retention capacity using as detailed previously^3,26–28^. Briefly, 2-4 million trypsinized cells, as indicated, were washed in fresh media, centrifuged for 5min at 200g, and resuspended in extracellular-like Ca^2+^-free buffer (120mM NaCl; 5mM KCl; 1mM KH_2_PO_4_; 0.2mM MgCl_2_ꞏ6H_2_O; 0.1mM EGTA; 20mM HEPES; pH 7.4), and incubated on ice for 5min to chelate extracellular Ca^2+^. Cells were centrifuged at 200g for 5min, the supernatant removed, and the pelleted cells resuspended in permeabilization buffer consisting of intracellular-like medium (120mM KCl; 10mM NaCl; 1mM KH_2_PO_4_; 20mM HEPES; pH 7.2) that had been cleared with Chelex 100 (Bio-Rad # 1422822) to remove trace Ca^2+^, and supplemented with 1X EDTA-free protease inhibitor cocktail (Sigma-Aldrich #4693132001), 80 µg/mL digitonin (Sigma-Aldrich # D141), 6µM thapsigargin (Enzo Life Sciences # BML-PE180-0005), and 5mM succinate (Sigma-Aldrich # S3674). 1 µM Fura-FF (AAT Bioquest #21028) was used to monitor extra-mitochondrial Ca^2+^ and 4.8 µM JC-1 (Enzo Life Sciences #52304) was used to monitor mitochondrial membrane potential (ΔΨ_m_) using a Quanta-Master-800 High Speed Ratiometric Spectrophotometer (Photon Technology International).

To measure _m_Ca^2+^ efflux, a 10µM bolus of CaCl_2_ was added at 350sec, followed by injection of 10µM Ru360 (EMD Millipore #557440-500UG), a mitochondrial calcium uniporter inhibitor, to assess _m_Ca^2+^ efflux independent of uptake. Experiments were terminated by injection of 10µM FCCP (Sigma-Aldrich #C2920) to collapse ΔΨ_m_ and release remaining free matrix Ca^2+^ to the bath solution. Where indicated, 20-40µM CGP-37157 (Enzo Life Sciences #BML-CM119-0005) was added from the beginning of the experiment to inhibit NCLX and evaluate NCLX-dependent _m_Ca^2+^ efflux. To measure mitochondrial calcium retention capacity, 5-10 µM boluses of CaCl_2_ were injected as indicated every 60 sec beginning at 400sec. CaCl_2_ injections continued until mitochondrial permeability transition, indicated by spontaneous collapse of ΔΨ_m_ and release of matrix Ca^2+^ to the bath solution. Experiments were terminated by addition of 10µM FCCP to confirm collapse of ΔΨ_m_.

### Live cell Ca^2+^ transient imaging

AC16 cardiomyocytes and *Nclx*^fl/fl^ MEFs were transduced with adenovirus encoding the mitochondrial Ca^2+^ reporter Mito-R-GECO 48 hours before imaging. Immediately prior to imaging, cells were loaded with the Ca^2+^-sensitive dye Fluo-4-AM (Thermo Fisher Scientific # F14201) at 1µM for 30min at 37°C, then washed twice in 37°C Tyrode’s buffer (150mM NaCl, 5.4mM KCl, 1.2mM MgCl_2_ꞏ6H_2_O, 10mM glucose, 2mM sodium pyruvate, 5mM HEPES, 1mM CaCl_2_ꞏ2H_2_O, pH = 7.4). Cells were imaged in fresh Tyrode’s buffer in a heated 37°C chamber using a Carl Zeiss Axio Observer Z1 microscope. Mitochondrial and cytosolic Ca^2+^ signals were continuously recorded at 572/35nm_ex_, 632/60nm_em_ and 490/20nm_ex_, 632/60nm_em_, respectively, and analyzed on Zen software (Zeiss). After 1min of baseline recording, a bolus of KCl was added to a final concentration of 100mM to depolarize the plasma membrane and allow influx of extracellular Ca^2+^. Mitochondrial and cytosolic Ca^2+^ transients were analyzed in Clampfit 10.7 software (Molecular Devices).

### Na^+^-stimulated _m_Ca^2+^ efflux

AC16 cardiomyocytes were trypsinized and counted. 100,000 cells were pelleted by centrifugation for 5min at 200g, then resuspended in 500µL Ca^2+^- and Na^+^-free extracellular-like buffer (120mM sucrose; 5mM KCl; 1mM KH_2_PO_4_; 0.2mM MgCl_2_ꞏ6H_2_O; 0.1mM EGTA; 20mM HEPES; pH 7.4), and incubated on ice for 5min to chelate extracellular Ca^2+^. Cells were centrifuged again, and the pellet resuspended in 37.5µL of Na^+^-free intracellular-like medium (120mM KCl; 1mM KH_2_PO_4_; 20mM HEPES; pH 7.2) that had been cleared with Chelex 100 (Bio-Rad # 1422822) to remove trace Ca^2+^, and supplemented with 80 µg/mL digitonin and 6µM thapsigargin. The 37.5µL cell suspension was transferred to a well of a 96-well plate (Greiner Bio-One # 655090) containing 12.5µL of assay buffer (125mM KCl, 20mM HEPES, 2mM MgCl_2_, 2mM KH_2_PO_4_, pH =7.4) supplemented with 40mM succinate (Sigma-Aldrich # S3674-100G), 40mM malate (Sigma-Aldrich #240176-50G), 40mM pyruvate (Sigma-Aldrich # P5280-100G), 4µM calcium green-5N hexapotassium salt (Invitrogen #C-3737) and, where indicated, 40µM CGP-37157 (final concentrations 10mM each succinate, malate, pyruvate; 1µM calcium green-5N; 10µM CGP-37157). Fluorescence was measured every 200ms at 506nm_ex_/532nm_em_ using a TECAN Infinite M1000 Pro plate reader set at 37°C. After 120sec of baseline recording, 5 successive 5µL injections of 10µM CaCl_2_ dissolved in assay buffer were administered at intervals of 120 seconds to increase the Ca^2+^ concentration by increments of 1µM. 120sec after the final Ca^2+^ bolus, NaCl dissolved in assay buffer was injected to a final concentration of 10mM to stimulate NCLX-dependent _m_Ca^2+^ efflux. _m_Ca^2+^ efflux rate was quantified over the first 75 seconds following the injection of NaCl.

### NCLX-3xFLAG mouse generation and validation

Single guide RNA (5’-GCTTCAGTCACGCCTTCTTC-3’) targeting exon 16 of the mouse *Slc8b1* gene, single-stranded oligodeoxynucleotide encoding an alanine linker and 3xFLAG epitope tag knocked in in-frame between the final coding codon and stop codon of mus *Slc8b1* (5’-catctgtttccttgttgtggtcctgctcacagagtttggggtgattcacctgaagaaggcagcggactacaaagaccatgacggtgattataaagatcatgacatcgattacaaggatgacgatgacaagtgactgaagctgcttggcctagaggtgtgggggcgattctgctagcctcctgagggggag-3’), and Cas9 protein were injected into the pronucleus of fertilized C57BL6/J ova, which were then transplanted into pseudo-pregnant female mice. After validation by sequencing and confirmation of germline transmission in founder lines, knock-in mice were back-crossed onto the C57BL/6NJ background to remove the *Nnt* mutation present in the C57BL6/J strain^29^.

Pups were genotyped for presence of the 3xFLAG knockin allele using the forward primer: 5’- ATCTTCTCCCTGGTCTCCGT-3’ and the reverse primer: 5’- GGACCTTCTCACCCATGCAG-3’. PCR reaction mixture contained 1uL tail DNA in DirectPCR Lysis reagent (Viagen Biotech #102- T), 1× Taq buffer (Syd Labs #MB042-EUT), 80 μM each dNTPs (New England Biolabs #N0447L), 800 nM each forward and reverse primers, and 1.25 U Taq polymerase (Syd Labs #MB042- EUT). The PCR conditions were: denaturation at 95°C for 3 min, followed by 35 cycles (95°C for 30 s, 60.5°C for 30 s, 72°C for 1min), followed by 10 min at 72°C. Pups were genotyped for the *Nnt* mutation using the WT forward primer: 5’- GGGCATAGGAAGCAAATACCAAGTTG-3’ and the mutant forward primer: 5’- GTGGAATTCCGCTGAGAGAACTCTT-3’ with the common reverse primer: 5’- GTAGGGCCAACTGTTTCTGCATGA-3’. PCR reaction mixture contained 1uL tail DNA in DirectPCR Lysis reagent, 1× Taq buffer, 80 μM each dNTPs, 800 nM each forward and reverse primers, and 1.25 U Taq polymerase. PCR conditions were: denaturation at 95°C for 3 min, followed by 35 cycles (95°C for 30 s, 60.5°C for 30 s, 72°C for 30sec, followed by 10 min at 72°C.

To verify mitochondrial expression of the NCLX-3xFLAG tagged protein, cardiac mitochondria were isolated from hearts of mice according to Frezza et al^30^. Following isolation, 500µg of mitochondria were progressively solubilized 1-4% digitonin (Sigma-Aldrich #D141-500MG) dissolved in water. Briefly, mitochondria were resuspended in 500µL of 1% digitonin, incubated on ice for 30min, then centrifuged at 12000g for 10min at 4°C. The supernatant was saved, and the resulting insoluble pellet was resuspended in 2% digitonin. The process was repeated, with the next pellet resuspended in 4% digitonin. Equal volumes of lysate supernatants in 1-4% digitonin were then used for western blotting.

### Fast protein size-exclusion liquid chromatography (FPLC)

Mitochondria isolated from AC16 cardiomyocytes or from 3-5 pooled mouse hearts were lysed on ice for 30min in 1x RIPA buffer with 1x SIGMAFAST protease inhibitor cocktail. Lysates were centrifuged at 14000g for 10min at 4°C and the protein concentration in the supernatant was determined by bicinchoninic acid assay. 2500µg of cleared mitochondrial lysate were fractionated by fast protein size-exclusion liquid chromatography (AKTA Pure FPLC; GE Healthcare), using a Superose 6 Increase 10/300 GL column (Sigma-Aldrich, # GE29-0915-96) equilibrated in 1X PBS, at a flow rate of 0.5mL/min. 0.5mL protein fractions were collected and concentrated to 75µL with 3kD molecular weight cutoff AMICON Ultra-0.5 centrifugal filter devices (EMD Millipore #UFC500396) according to the manufacturer’s instructions. Concentrated fractions #7-31 were used for western blotting. Molecular weights of FPLC fractions were calibrated using gel filtration standards (Bio-Rad # 1511901) and fitting their elution pattern to a one-phase exponential decay.

### *In silico* modeling

Models of the human NCLX (UniProt #Q6J4K2) binding to human TMEM65 (UniProt #Q6PI78) proteins were generated as 1:1 heterodimers with AlphaFold2 implemented in ColabFold^31^ using the advanced workbook and various options (MMseqs2, unpaired, minimum coverage of 75%). The resulting models were ranked by pTMscore and visualized in UCSF Chimera^32^ and ICM-Browser Pro (MolSoft).

### *In vivo* knockdown of *Tmem65*

Male and female C57BL/6NJ mice at postnatal day 1-3 were administered 2.25^10^11 viral particles of adeno-associated virus 9 (AAV9) via intracardiac injection. Viruses included AAV9 encoding scrambled shRNA (5’-CCTAAGGTTAAGTCGCCCTCG-3’) under the control of a U6 promoter, and AAV9 encoding shRNA against murine *Tmem65* (5’- ACTATGGAAACACTAACTTAT-3’) under the control of a U6 promoter. End-point assessments for cardiac muscle function and pathology were conducted at 6 weeks of age, and end-point assessments for neuromuscular function and skeletal muscle pathology were conducted at 24 weeks of age. All *in vivo* experiments and analyses were performed using a numbered ear-tagging system in order to blind the experimenter to mouse genotype and experimental group. All animal experiments followed AAALAC guidelines and were approved by Temple University’s IACUC.

### Four-limb hanging test

To assess integrated neuromuscular function, mice were placed on a wire grid. After 30 sec of acclimation, the grid was inverted at a height of 35mm over soft bedding. The time until the mice fell off of the inverted grid was recorded as hang time. The procedure was repeated 3 times with a rest interval of at least 10 minutes between trials, and the mean hang time for each mouse was used to calculate its mean holding impulse in N/s (body weight (g) x mean hang time (s) x 9.806^10^-3N/g), according to established protocols (TREAT-NMD Neuromuscular Network, SOP # DMD_M.2.1.005 and^33^).

### Left ventricular echocardiography

Mice were anesthetized and transthoracic echocardiography of the left ventricle was performed and analyzed as described previously^5^.

### Tissue gravimetrics and histology

Animals were euthanized at the indicated endpoints. Tibia length and total body mass were recorded for normalization of heart and skeletal muscle mass. The heart base was snap frozen in liquid nitrogen then processed for western blotting as detailed elsewhere^5^. The gastrocnemius muscle was dissected from one leg and snap frozen in liquid nitrogen prior to processing for western blotting. The gastrocnemius + soleus + plantaris muscles were dissected from the other leg as unit and massed, then placed in Tissue-Tek O.C.T. compound (Sakura #4583) and frozen in liquid nitrogen-cooled isopentane. Muscles were cut to 8µm cross sections, mounted on glass slides, and labelled with wheat germ agglutinin-AlexaFluor 488 conguate (ThermoFisher #W11261) and DAPI as described^28^. Slides were imaged at 10x magnification on a Nikon Eclipse Ti-E fluorescence microscope. Muscle fiber cross sectional area, minimal Feret’s diameter, and the percentage of centrally nucleated fibers were measured for at least 150 fibers per muscle in ImageJ (National Institutes of Health). The variance coefficient of the minimal Feret’s diameter was calculated as: (standard deviation of minimal Feret’s diameter / mean minimal Feret’s diameter) x 100^34^.

### Imaging of mitochondrial membrane potential, ΔΨ_m_

Cells were cultured on 35mm glass-bottomed dishes and prior to imaging, incubated in Tyrode’s buffer containing 15nM tetramethylrhodamine, methyl ester (TMRM) (Thermo Fisher Scientific #T668) at 37°C for 15min. Cells were then washed 2x in fresh 37°C Tyrode’s buffer, and changed to 2mL fresh Tyrode’s buffer supplemented with either 10µM cyclosporine A (CsA) (Sigma-Aldrich #30024-25MG) or ethanol as the vehicle control and incubated at 37°C. After 10 minutes, cells were imaged at 545/30nm_ex_/ 585/40nm_em_ in a heated 37°C chamber using a Carl Zeiss Axio Observer Z1 microscope. Images were acquired for 5-6 20x fields of view, and TMRM staining intensity quantified with Zen software. Specificity of the TMRM signal was confirmed at the end of imaging by addition of FCCP to a final concentration of 10µM to depolarize mitochondria.

### *In vitro* cell death assay

Cells were plated in 96-well plates at 500cells/well. The following day, 0-20µM thapsigargin or ionomycin (Cayman Chemical #11932) was added to the culture media to induce cellular Ca^2+^ stress. After 24 hours, Sytox Green (Invitrogen #S7020) was added at a final concentration of 5µM to assess plasma membrane rupture. One well per plate had dye omitted as a control for background fluorescence. The cells were incubated for a further 30min at 37°C, then Sytox Green fluorescence was measured at 504nm_ex/_523nm_em_ on a TECAN Infinite M1000 Pro plate reader. Hoechst 33342 (Thermo Fisher Scientific #62249) was then added at a final concentration of 10µM and the cells were incubated for another 10 min at 37°C. One well per plate had dye omitted as a control for background fluorescence. Hoechst fluorescence was measured with the plate reader at 350nm_ex_/461nm_em_. Sytox green fluorescence was normalized to Hoechst fluorescence within each well to account for differences in total cell number.

### Statistical analysis

All results are presented as mean ± S.E.M unless otherwise noted. Statistical analysis was performed using Prism 6.0 (GraphPad Software). Direct comparisons between two groups used an unpaired two-tailed t-test, with Welch’s correction cases of unequal variance. Where data did not follow a normal distribution, a non-parametric Mann-Whitney test was used. Simple comparisons between more than two groups used 1-way ANOVA followed by Dunnet’s post-hoc test for comparisons to a single control group or by Tukey’s post-hoc test for comparisons among all groups. Grouped data were analyzed by 2-way ANOVA with Sidak’s post-hoc analysis. Correlation between variables was evaluated by linear regression. For all comparisons, *P* values less than 0.05 were considered significant.

### Data availability

All relevant data are available from the authors. The mass spectrometry proteomics data for the proximity biotinylation screen were uploaded to the ProteomeXchange Consortium via MassIVE with dataset identifiers PXD045834 and MSV000093004, respectively.

## Abbreviations

Ca^2+^: calcium
ΔΨ_m_: mitochondrial membrane potential
IMM: inner mitochondrial membrane
_m_Ca^2+^: mitochondrial calcium
Na^+^: sodium
NCLX: mitochondrial sodium-calcium exchanger

## Acknowledgments

We thank Xiaoxia Cui and the Genome Engineering and iPSC Center (GEiC) at Washington University in St. Louis for gRNA validation and genotyping services; Xiang Hua and the Fox Chase Cancer Center Transgenic Mouse Facility for assistance in generating the NCLX- 3xFLAG mouse line; and Mark Andrake of the Fox Chase Cancer Center Molecular Modeling Facility for assistance with *in silico* modeling of protein-protein interactions.

The research was supported by the NIH (T32HL091804 and F32HL151146 to J.F.G..; T32HL091804 and F30AG082407 to H.M.C.; DK120876 to D.T.; R01NS121379, P01HL147841, P01HL134608, R01HL142271 to J.W.E.), a Frances Velay Fellowship (to A.E.S.), and the American Heart Association (20EIA35320226 to J.W.E.).

## Author Contributions

Conception and design of research: J.F.G. and J.W.E.

Performed experiments: J.F.G., O.S., H.M.C. C.C-F., A.S.M, A.D.M., A.E.S., D.Y.K, J.J.D., A.W., E.K.M., M.P.L., A.N.H., and D.T.

Analyzed data: J.F.G., H.M.C., C.C-F., D.T., and J.W.E.

Interpreted results of experiments: J.F.G. and J.W.E.

Prepared figures: J.F.G.

Drafted manuscript: J.F.G.

Edited and revised manuscript: J.F.G. and J.W.E.

Approved final version of manuscript: J.F.G., O.S., H.M.C. C.C-F., A.S.M, A.D.M., A.E.S., D.Y.K, J.J.D., A.W., E.K.M., M.P.L., A.N.H., D.T., and J.W.E.

## Materials and Correspondence

Correspondence and requests for materials should be addressed to J.W.E.

---

**Supplemental Fig. 1:**
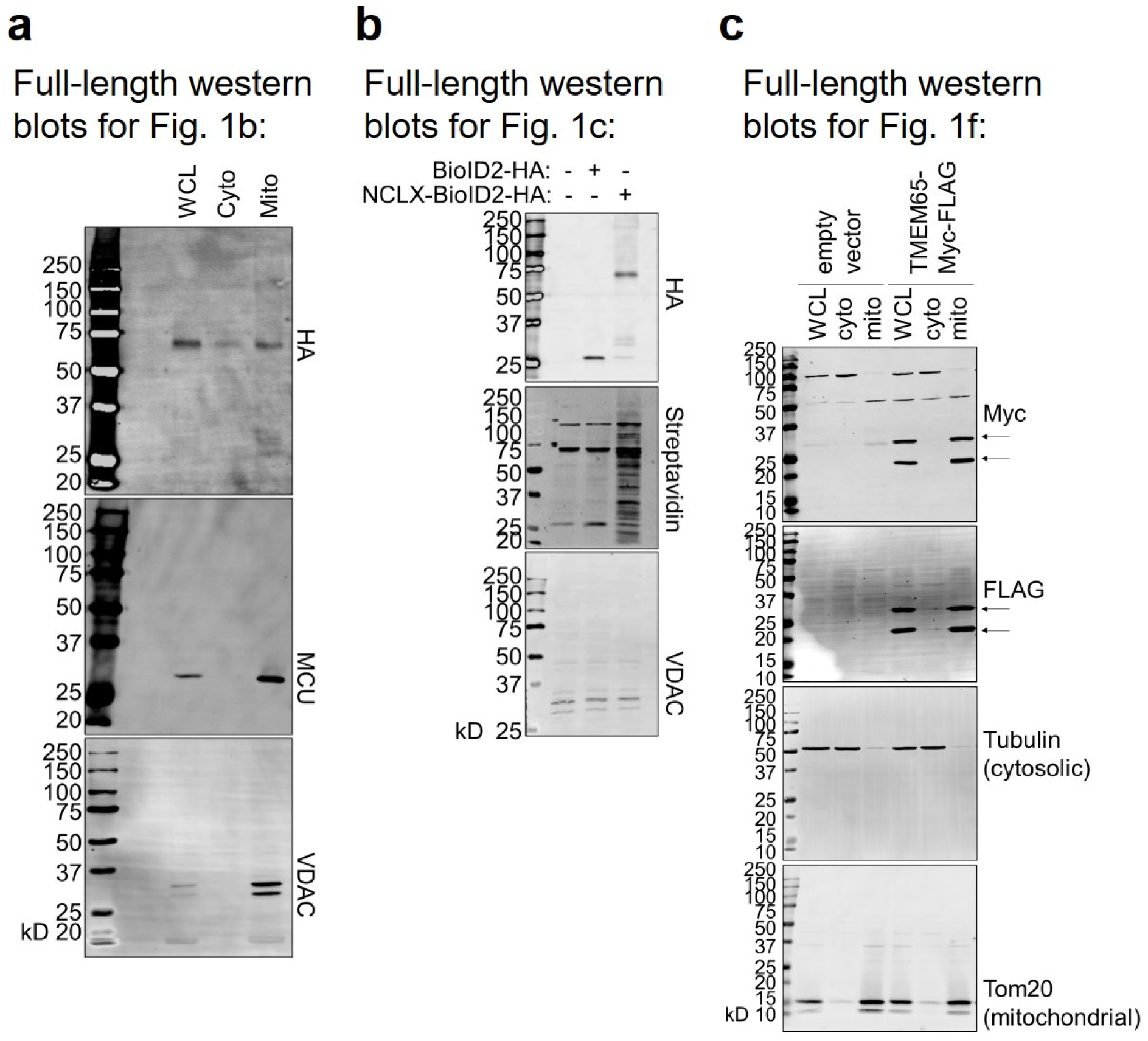
Full-length western blots for Fig. 1.

**Supplemental Fig. 2:**
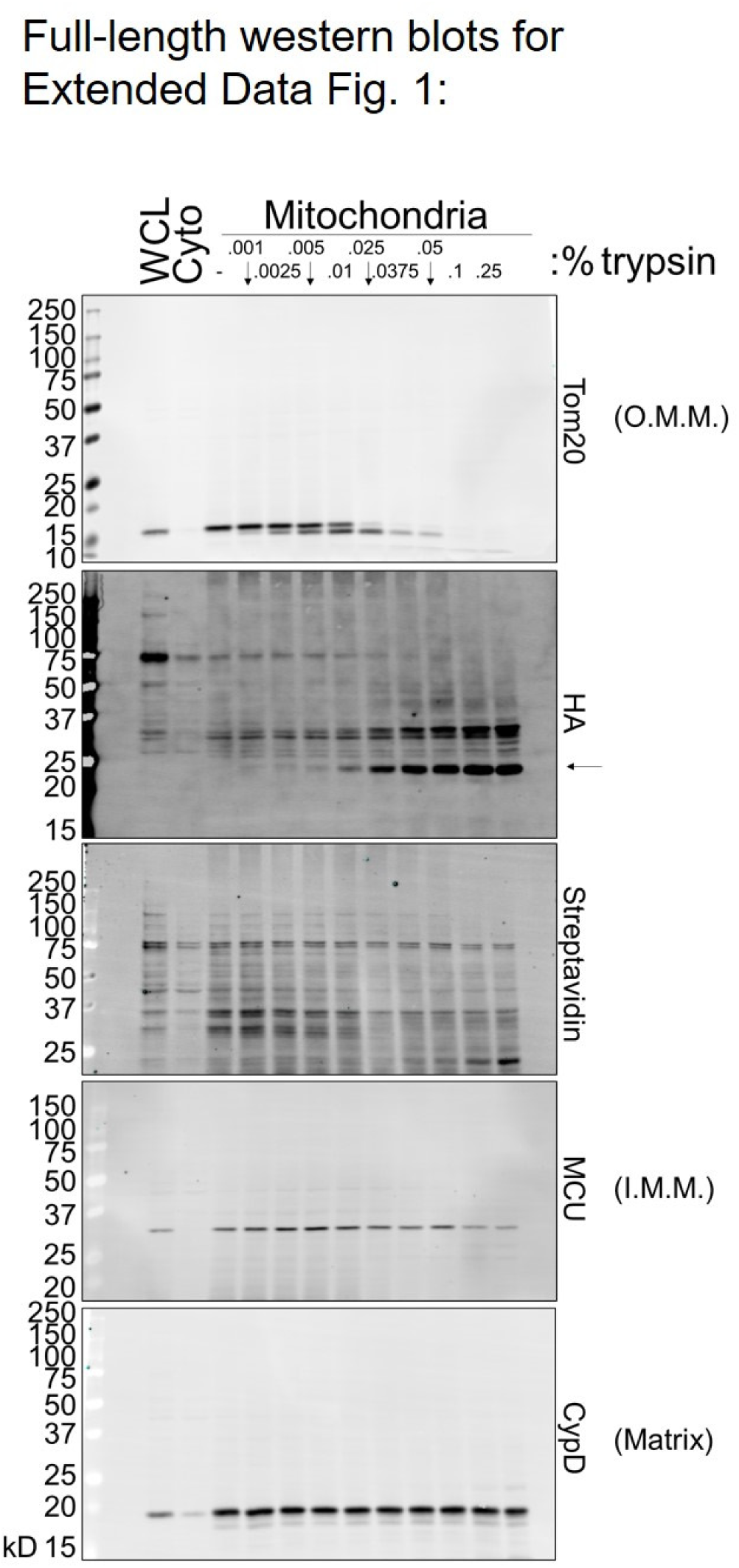
Full-length western blots for Extended Data Fig. 1.

**Supplemental Fig. 3:**
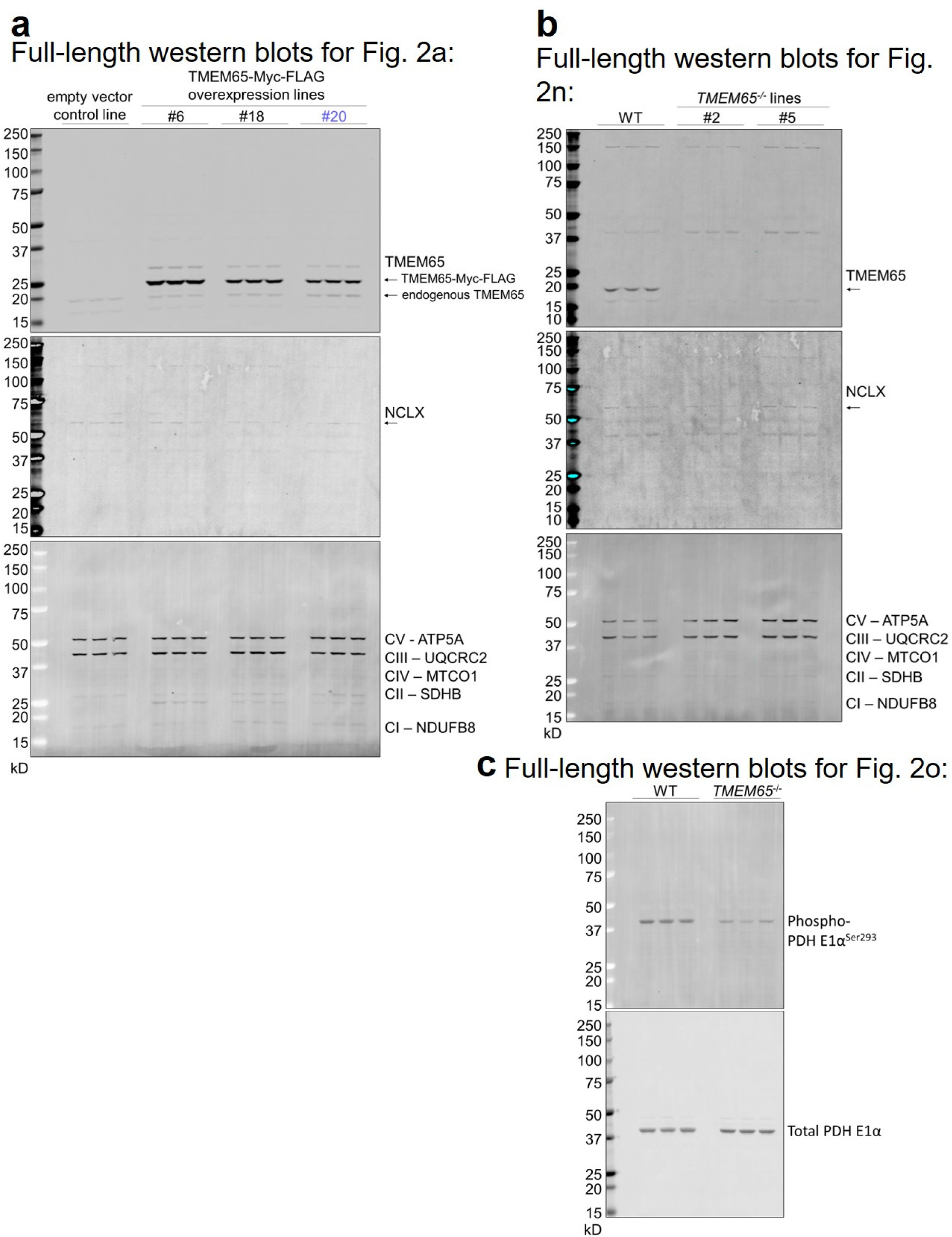
Full-length western blots for Fig. 2.

**Supplemental Fig. 4:**
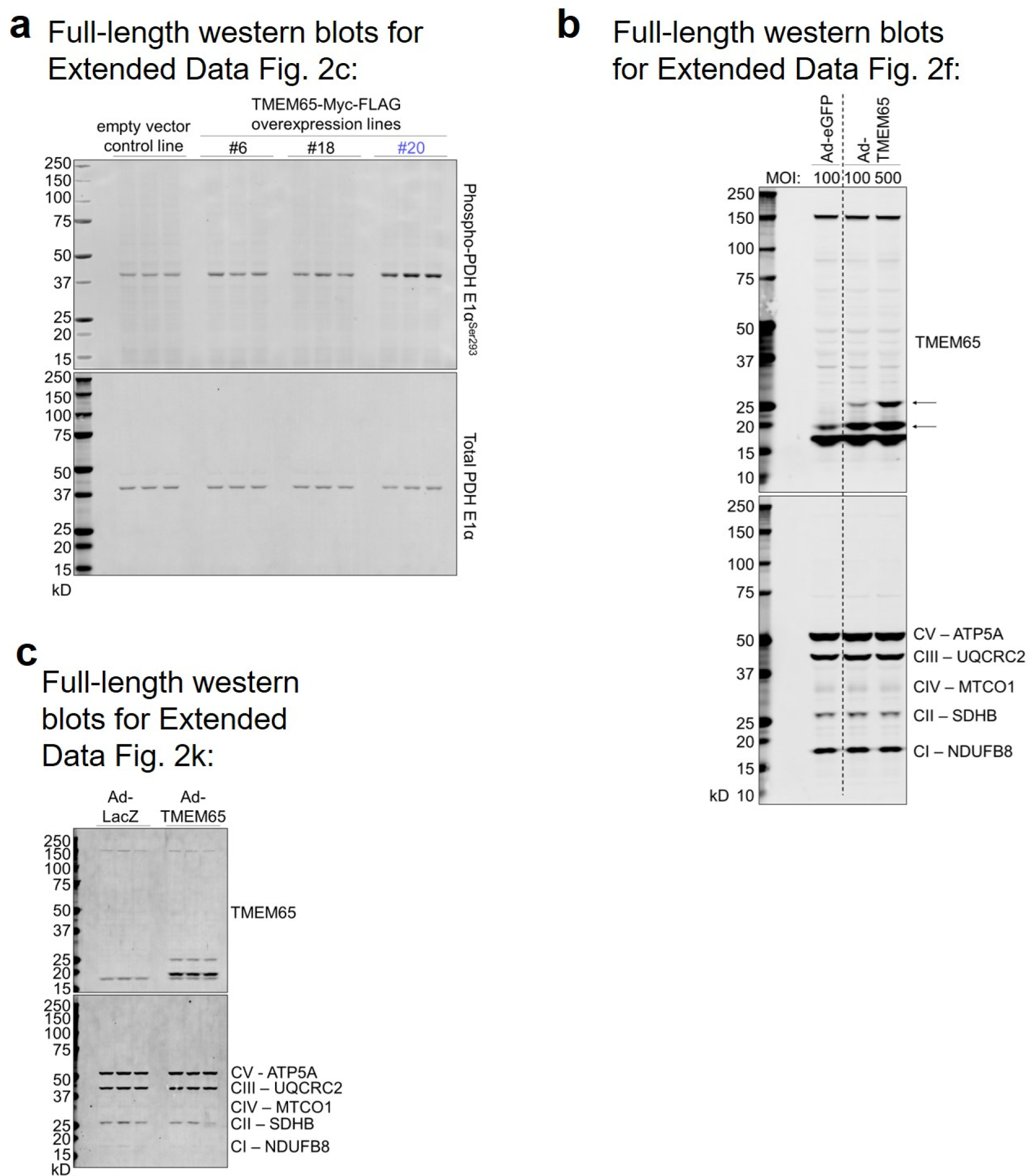
Full-length blots for Extended Data Fig. 2.

**Supplemental Fig. 5:**
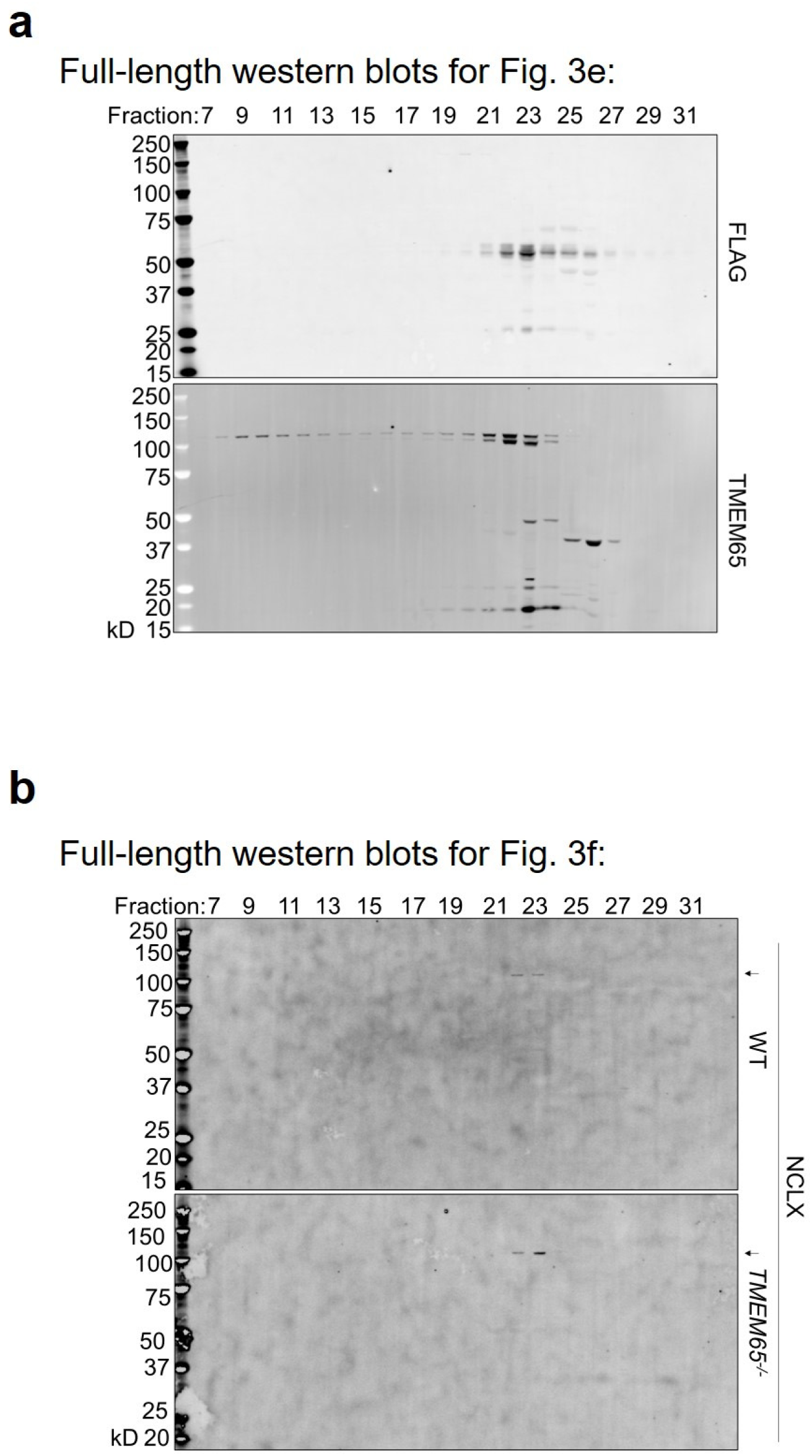
Full-length western blots for Fig. 3.

**Supplemental Fig. 6:**
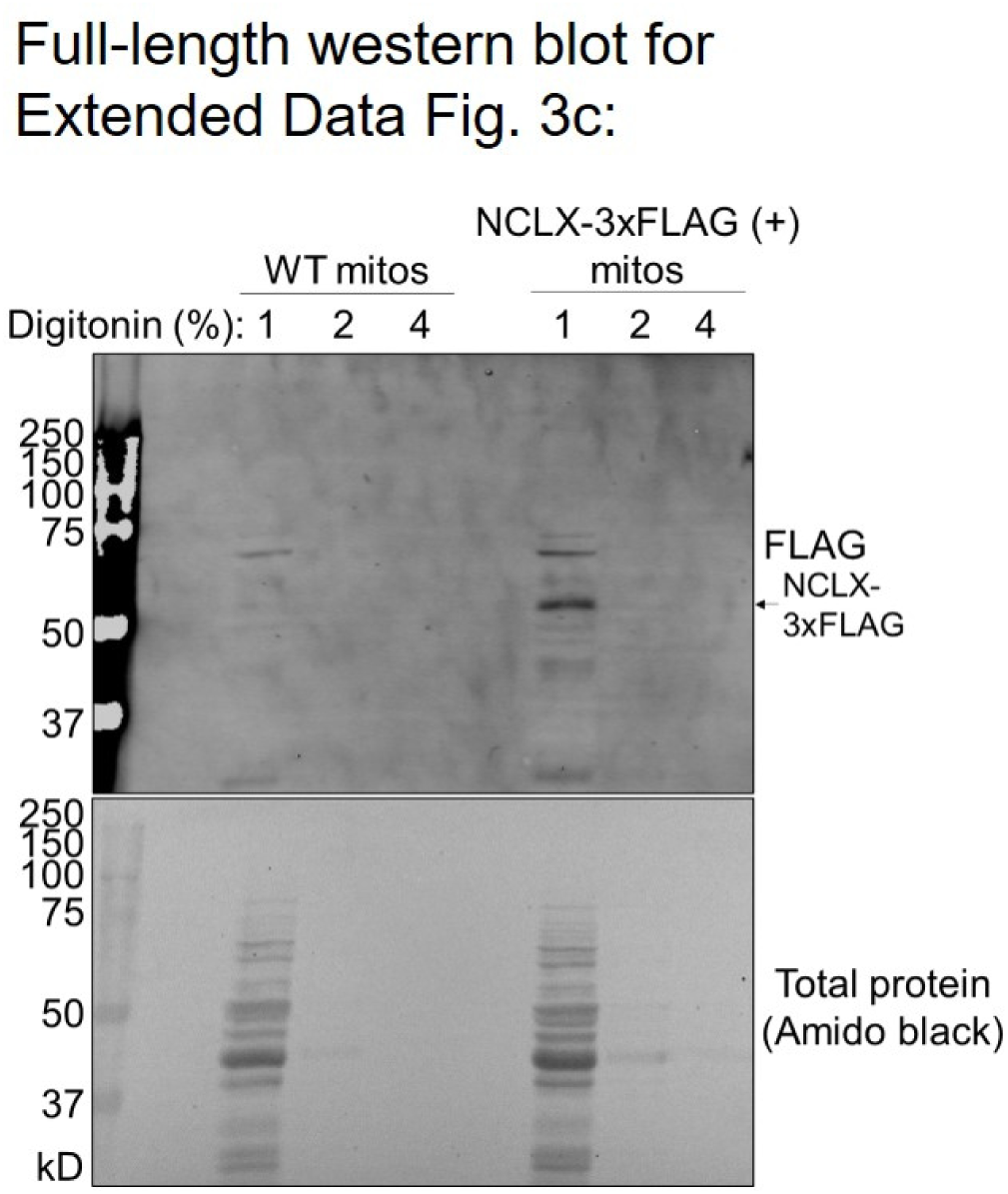
Full-length western blot for Extended Data Fig. 3.

**Supplemental Fig. 7:**
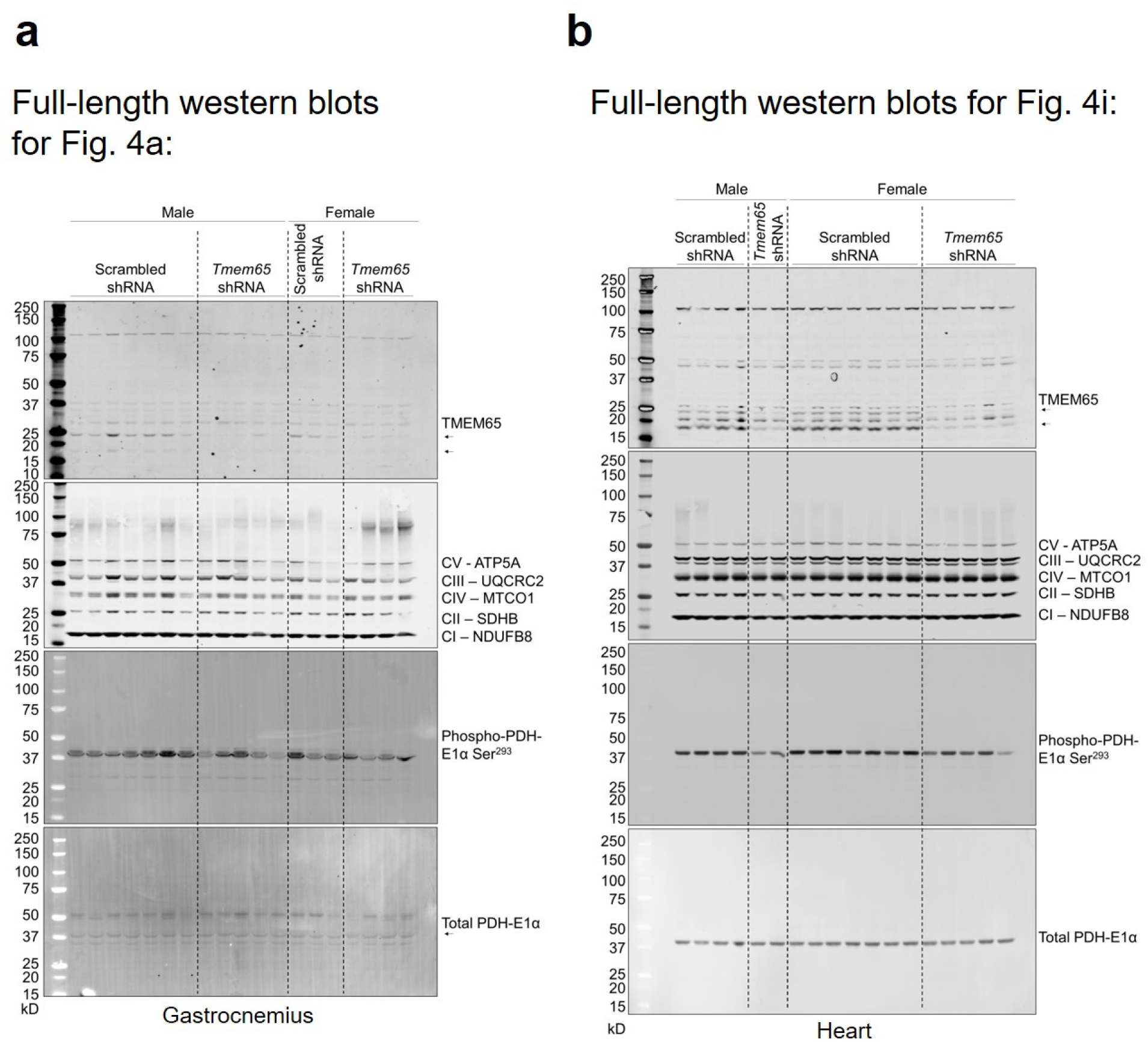
Full-length western blots for Fig. 4.

**<u>Tables:</u>**

**Extended Data Table 1:** Known and predicted mitochondrial proteins enriched ≥2x in NCLX-BioID2-HA over BioID2-HA control in ≥2 out of 3 replicates.

