## Extended Data Table 1 for "TMEM65 regulates NCLX-dependent mitochondrial calcium efflux"

**Extended Data Table 1: Known and predicted mitochondrial proteins enriched ≥2x in NCLX-BioID2-HA over BioID2-HA control in ≥2 out of 3 replicates**

| Protein IDs | Majority protein IDs | Protein names | Gene names | Fasta headers | LFQ intensity AC16_Ctrl_RepA | LFQ intensity AC16_Ctrl_RepB | LFQ intensity AC16_Ctrl_RepC | LFQ intensity AC16_hNCLX_RepA | LFQ intensity AC16_hNCLX_RepB | LFQ intensity AC16_hNCLX_RepC | Fold change LFQ AC16_hNCLX/LFQ AC16_Ctrl_RepA | Fold change LFQ AC16_hNCLX/LFQ AC16_Ctrl_RepB | Fold change LFQ AC16_hNCLX/LFQ AC16_Ctrl_RepC |
|---|---|---|---|---|---|---|---|---|---|---|---|---|---|
| P27824 | P27824 | Calnexin | CANX | Calnexin OS=Homo sapiens OX=9606 GN=CANX PE=1 SV=2 | 2.139E+09 | 0 | 785430000 | 6.987E+09 | 5.711E+09 | 1.408E+09 | 3.2659281 | #DIV/0! | 1.7929033 |
| Q6J4K2 | Q6J4K2 | Sodium/potassium/calcium exchanger 6, mitochondrial | SLC8B1 | Mitochondrial sodium/calcium exchanger protein OS=Homo sapiens OX=9606 GN=SLC8B1 PE=1 SV=2 | 0 | 0 | 0 | 1.597E+09 | 2.297E+09 | 161370000 | #DIV/0! | #DIV/0! | #DIV/0! |
| Q969Z0 | Q969Z0 | Protein TBRG4 | TBRG4 | FAST kinase domain-containing protein 4 OS=Homo sapiens OX=9606 GN=TBRG4 PE=1 SV=1 | 0 | 0 | 0 | 71443000 | 86241000 | 22803000 | #DIV/0! | #DIV/0! | #DIV/0! |
| Q9Y673 | Q9Y673 | Dolichyl-phosphate beta-glucosyltransferase | ALG5 | Dolichyl-phosphate beta-glucosyltransferase OS=Homo sapiens OX=9606 GN=ALG5 PE=1 SV=1 | 0 | 0 | 0 | 74919000 | 67498000 | 26056000 | #DIV/0! | #DIV/0! | #DIV/0! |
| P35580 | P35580 | Myosin-10 | MYH10 | Myosin-10 OS=Homo sapiens OX=9606 GN=MYH10 PE=1 SV=3 | 15193000 | 0 | 11104000 | 98862000 | 154050000 | 32285000 | 6.5070756 | #DIV/0! | 2.9075108 |
| A1L0T0 | A1L0T0 | Acetolactate synthase-like protein | ILVBL | Acetolactate synthase-like protein OS=Homo sapiens OX=9606 GN=ILVBL PE=1 SV=2 | 56762000 | 0 | 7766900 | 115060000 | 136090000 | 21902000 | 2.0270604 | #DIV/0! | 2.8199153 |
| Q9NSE4 | Q9NSE4 | Isoleucine--tRNA ligase, mitochondrial | IARS2 | Isoleucine--tRNA ligase, mitochondrial OS=Homo sapiens OX=9606 GN=IARS2 PE=1 SV=2 | 48604000 | 0 | 23009000 | 116350000 | 135740000 | 11144000 | 2.3938359 | #DIV/0! | 0.4843322 |
| Q8NE86 | Q8NE86 | Calcium uniporter protein, mitochondrial | MCU | Calcium uniporter protein, mitochondrial OS=Homo sapiens OX=9606 GN=MCU PE=1 SV=1 | 0 | 0 | 0 | 50964000 | 63164000 | 18717000 | #DIV/0! | #DIV/0! | #DIV/0! |
| P00167 | P00167 | Cytochrome b5 | CYB5A | Cytochrome b5 OS=Homo sapiens OX=9606 GN=CYB5A PE=1 SV=2 | 0 | 0 | 0 | 68290000 | 41261000 | 18736000 | #DIV/0! | #DIV/0! | #DIV/0! |
| P50416;Q92523 | P50416 | Carnitine O-palmitoyltransferase 1, liver isoform | CPT1A | Carnitine O-palmitoyltransferase 1, liver isoform OS=Homo sapiens OX=9606 GN=CPT1A PE=1 SV=2 | 0 | 0 | 0 | 76117000 | 38974000 | 0 | #DIV/0! | #DIV/0! | #DIV/0! |
| O43852 | O43852 | Calumenin | CALU | Calumenin OS=Homo sapiens OX=9606 GN=CALU PE=1 SV=2 | 0 | 0 | 0 | 48612000 | 39369000 | 26106000 | #DIV/0! | #DIV/0! | #DIV/0! |
| Q96EY7 | Q96EY7 | Pentatricopeptide repeat domain-containing protein 3, mitochondrial | PTCD3 | Pentatricopeptide repeat domain-containing protein 3, mitochondrial OS=Homo sapiens OX=9606 GN=PTCD3 PE=1 SV=3 | 0 | 0 | 0 | 31065000 | 43722000 | 38152000 | #DIV/0! | #DIV/0! | #DIV/0! |
| Q9BYK8 | Q9BYK8 | Helicase with zinc finger domain 2 | HELZ2 | Helicase with zinc finger domain 2 OS=Homo sapiens OX=9606 GN=HELZ2 PE=1 SV=6 | 0 | 0 | 0 | 35346000 | 75968000 | 0 | #DIV/0! | #DIV/0! | #DIV/0! |
| Q96GF1 | Q96GF1 | E3 ubiquitin-protein ligase RNF185 | RNF185 | E3 ubiquitin-protein ligase RNF185 OS=Homo sapiens OX=9606 GN=RNF185 PE=1 SV=1 | 0 | 0 | 0 | 56598000 | 51372000 | 0 | #DIV/0! | #DIV/0! | #DIV/0! |
| Q6IAN0 | Q6IAN0 | Dehydrogenase/reductase SDR family member 7B | DHRS7B | Dehydrogenase/reductase SDR family member 7B OS=Homo sapiens OX=9606 GN=DHRS7B PE=1 SV=2 | 0 | 0 | 0 | 59546000 | 39730000 | 6842200 | #DIV/0! | #DIV/0! | #DIV/0! |
| O75153;Q9Y5I4 | O75153 | Clustered mitochondria protein homolog | CLUH | Clustered mitochondria protein homolog OS=Homo sapiens OX=9606 GN=CLUH PE=1 SV=2 | 0 | 24797000 | 8537000 | 81797000 | 153700000 | 13507000 | #DIV/0! | 6.1983304 | 1.5821717 |
| O95870 | O95870 | Abhydrolase domain-containing protein 16A | ABHD16A | Phosphatidylserine lipase ABHD16A OS=Homo sapiens OX=9606 GN=ABHD16A PE=1 SV=3 | 0 | 0 | 0 | 36232000 | 37876000 | 5443200 | #DIV/0! | #DIV/0! | #DIV/0! |
| P32119 | P32119 | Peroxiredoxin-2 | PRDX2 | Peroxiredoxin-2 OS=Homo sapiens OX=9606 GN=PRDX2 PE=1 SV=5 | 0 | 0 | 0 | 40255000 | 18862000 | 14119000 | #DIV/0! | #DIV/0! | #DIV/0! |
| P17152 | P17152 | Transmembrane protein 11, mitochondrial | TMEM11 | Transmembrane protein 11, mitochondrial OS=Homo sapiens OX=9606 GN=TMEM11 PE=1 SV=1 | 0 | 0 | 0 | 44463000 | 28217000 | 0 | #DIV/0! | #DIV/0! | #DIV/0! |
| Q8NBU5 | Q8NBU5 | ATPase family AAA domain-containing protein 1 | ATAD1 | ATPase family AAA domain-containing protein 1 OS=Homo sapiens OX=9606 GN=ATAD1 PE=1 SV=1 | 0 | 0 | 0 | 42974000 | 23649000 | 0 | #DIV/0! | #DIV/0! | #DIV/0! |
| Q12981 | Q12981 | Vesicle transport protein SEC20 | BNIP1 | Vesicle transport protein SEC20 OS=Homo sapiens OX=9606 GN=BNIP1 PE=1 SV=3 | 0 | 0 | 0 | 37105000 | 26889000 | 0 | #DIV/0! | #DIV/0! | #DIV/0! |
| O95721 | O95721 | Synaptosomal-associated protein 29 | SNAP29 | Synaptosomal-associated protein 29 OS=Homo sapiens OX=9606 GN=SNAP29 PE=1 SV=1 | 0 | 0 | 0 | 29371000 | 24083000 | 9445000 | #DIV/0! | #DIV/0! | #DIV/0! |

| | | | | | | | | | | | | | |
|---|---|---|---|---|---|---|---|---|---|---|---|---|---|
| Q8WUH6 | Q8WUH6 | Transmembrane protein 263 | TMEM263 | Transmembrane protein 263 OS=Homo sapiens OX=9606 GN=TMEM263 PE=1 SV=1 | 0 | 34760000 | 0 | 41857000 | 40757000 | 20977000 | #DIV/0! | 1.1725259 | #DIV/0! |
| Q9BXK5 | Q9BXK5 | Bcl-2-like protein 13 | BCL2L13 | Bcl-2-like protein 13 OS=Homo sapiens OX=9606 GN=BCL2L13 PE=1 SV=1 | 0 | 0 | 0 | 33046000 | 28211000 | 0 | #DIV/0! | #DIV/0! | #DIV/0! |
| Q8N0X7 | Q8N0X7 | Spartin | SPG20 | Spartin OS=Homo sapiens OX=9606 GN=SPART PE=1 SV=1 | 0 | 0 | 0 | 11339000 | 0 | 49454000 | #DIV/0! | #DIV/0! | #DIV/0! |
| Q9GZY8 | Q9GZY8 | Mitochondrial fission factor | MFF | Mitochondrial fission factor OS=Homo sapiens OX=9606 GN=MFF PE=1 SV=1 | 0 | 0 | 0 | 27928000 | 22514000 | 10076000 | #DIV/0! | #DIV/0! | #DIV/0! |
| O15270 | O15270 | Serine palmitoyltransferase 2 | SPTLC2 | Serine palmitoyltransferase 2 OS=Homo sapiens OX=9606 GN=SPTLC2 PE=1 SV=1 | 0 | 0 | 0 | 31325000 | 28768000 | 0 | #DIV/0! | #DIV/0! | #DIV/0! |
| Q8IYB8 | Q8IYB8 | ATP-dependent RNA helicase SUPV3L1, mitochondrial | SUPV3L1 | ATP-dependent RNA helicase SUPV3L1, mitochondrial OS=Homo sapiens OX=9606 GN=SUPV3L1 PE=1 SV=1 | 0 | 0 | 0 | 14435000 | 13135000 | 32457000 | #DIV/0! | #DIV/0! | #DIV/0! |
| Q9NX47 | Q9NX47 | E3 ubiquitin-protein ligase MARCH5 | MARCH5 | E3 ubiquitin-protein ligase MARCH5 OS=Homo sapiens OX=9606 GN=MARCH5 PE=1 SV=1 | 38200000 | 0 | 0 | 52252000 | 51163000 | 8727800 | 1.3678534 | #DIV/0! | #DIV/0! |
| Q96K37 | Q96K37 | Solute carrier family 35 member E1 | SLC35E1 | Solute carrier family 35 member E1 OS=Homo sapiens OX=9606 GN=SLC35E1 PE=1 SV=2 | 0 | 0 | 0 | 26189000 | 23430000 | 10200000 | #DIV/0! | #DIV/0! | #DIV/0! |
| Q9BQA9 | Q9BQA9 | Uncharacterized protein C17orf62 | C17orf62 | Cytochrome b-245 chaperone 1 OS=Homo sapiens OX=9606 GN=CYBC1 PE=1 SV=1 | 0 | 0 | 0 | 31955000 | 22567000 | 0 | #DIV/0! | #DIV/0! | #DIV/0! |
| Q9H5Q4 | Q9H5Q4 | Dimethyladenosine transferase 2, mitochondrial | TFB2M | Dimethyladenosine transferase 2, mitochondrial OS=Homo sapiens OX=9606 GN=TFB2M PE=1 SV=1 | 0 | 0 | 8381500 | 26930000 | 26424000 | 10930000 | #DIV/0! | #DIV/0! | 1.3040625 |
| Q8IXI1 | Q8IXI1 | Mitochondrial Rho GTPase 2 | RHOT2 | Mitochondrial Rho GTPase 2 OS=Homo sapiens OX=9606 GN=RHOT2 PE=1 SV=2 | 28196000 | 0 | 0 | 52265000 | 38113000 | 14525000 | 1.8536317 | #DIV/0! | #DIV/0! |
| Q9H845 | Q9H845 | Acyl-CoA dehydrogenase family member 9, mitochondrial | ACAD9 | Complex I assembly factor ACAD9, mitochondrial OS=Homo sapiens OX=9606 GN=ACAD9 PE=1 SV=1 | 24870000 | 0 | 24555000 | 64482000 | 52051000 | 34158000 | 2.5927624 | #DIV/0! | 1.3910812 |
| Q13617 | Q13617 | Cullin-2 | CUL2 | Cullin-2 OS=Homo sapiens OX=9606 GN=CUL2 PE=1 SV=2 | 0 | 30164000 | 0 | 51667000 | 73051000 | 0 | #DIV/0! | 2.4217942 | #DIV/0! |
| P24752 | P24752 | Acetyl-CoA acetyltransferase, mitochondrial | ACAT1 | Acetyl-CoA acetyltransferase, mitochondrial OS=Homo sapiens OX=9606 GN=ACAT1 PE=1 SV=1 | 0 | 0 | 0 | 0 | 6558400 | 44676000 | #DIV/0! | #DIV/0! | #DIV/0! |
| Q9NYY8 | Q9NYY8 | FAST kinase domain-containing protein 2 | FASTKD2 | FAST kinase domain-containing protein 2, mitochondrial OS=Homo sapiens OX=9606 GN=FASTKD2 PE=1 SV=1 | 0 | 0 | 0 | 14093000 | 30939000 | 4687300 | #DIV/0! | #DIV/0! | #DIV/0! |
| P22061 | P22061 | Protein-L-isoaspartate(D-aspartate) O-methyltransferase | PCMT1 | Protein-L-isoaspartate(D-aspartate) O-methyltransferase OS=Homo sapiens OX=9606 GN=PCMT1 PE=1 SV=4 | 0 | 0 | 0 | 15000000 | 20216000 | 13236000 | #DIV/0! | #DIV/0! | #DIV/0! |
| O75027 | O75027 | ATP-binding cassette sub-family B member 7, mitochondrial | ABCB7 | ATP-binding cassette sub-family B member 7, mitochondrial OS=Homo sapiens OX=9606 GN=ABCB7 PE=1 SV=2 | 16956000 | 0 | 0 | 37865000 | 33427000 | 13619000 | 2.2331328 | #DIV/0! | #DIV/0! |
| Q8WVC6 | Q8WVC6 | Dephospho-CoA kinase domain-containing protein | DCAKD | Dephospho-CoA kinase domain-containing protein OS=Homo sapiens OX=9606 GN=DCAKD PE=1 SV=1 | 0 | 0 | 4916500 | 25158000 | 18630000 | 8909300 | #DIV/0! | #DIV/0! | 1.8121224 |
| Q9BXW7 | Q9BXW7 | Cat eye syndrome critical region protein 5 | CECR5 | Haloacid dehalogenase-like hydrolase domain-containing 5 OS=Homo sapiens OX=9606 GN=HDHD5 PE=1 SV=1 | 0 | 0 | 0 | 16149000 | 22418000 | 4558500 | #DIV/0! | #DIV/0! | #DIV/0! |
| O15091 | O15091 | Mitochondrial ribonuclease P protein 3 | KIAA0391 | Mitochondrial ribonuclease P catalytic subunit OS=Homo sapiens OX=9606 GN=PRORP PE=1 SV=2 | 0 | 0 | 0 | 20595000 | 0 | 20166000 | #DIV/0! | #DIV/0! | #DIV/0! |
| Q92552 | Q92552 | 28S ribosomal protein S27, mitochondrial | MRPS27 | 28S ribosomal protein S27, mitochondrial OS=Homo sapiens OX=9606 GN=MRPS27 PE=1 SV=3 | 31745000 | 13142000 | 0 | 97341000 | 68456000 | 39041000 | 3.0663412 | 5.2089484 | #DIV/0! |
| Q9HD20 | Q9HD20 | Manganese-transporting ATPase 13A1 | ATP13A1 | Manganese-transporting ATPase 13A1 OS=Homo sapiens OX=9606 GN=ATP13A1 PE=1 SV=2 | 55400000 | 10004000 | 0 | 254200000 | 215450000 | 38175000 | 4.5884477 | 21.536385 | #DIV/0! |
| Q6AI08 | Q6AI08 | HEAT repeat-containing protein 6 | HEATR6 | HEAT repeat-containing protein 6 OS=Homo sapiens OX=9606 GN=HEATR6 PE=1 SV=1 | 0 | 0 | 0 | 8714000 | 26690000 | 0 | #DIV/0! | #DIV/0! | #DIV/0! |
| Q15388 | Q15388 | Mitochondrial import receptor subunit TOM20 homolog | TOMM20 | Mitochondrial import receptor subunit TOM20 homolog OS=Homo sapiens OX=9606 GN=TOMM20 PE=1 SV=1 | 0 | 0 | 0 | 5185700 | 0 | 30089000 | #DIV/0! | #DIV/0! | #DIV/0! |

| | | | | | | | | | | | | | |
|---|---|---|---|---|---|---|---|---|---|---|---|---|---|
| Q9NVV4 | Q9NVV4 | Poly(A) RNA polymerase, mitochondrial | MTPAP | Poly(A) RNA polymerase, mitochondrial OS=Homo sapiens OX=9606 GN=MTPAP PE=1 SV=1 | 0 | 0 | 0 | 14278000 | 20191000 | 0 | #DIV/0! | #DIV/0! | #DIV/0! |
| Q9Y5Z9 | Q9Y5Z9 | UbiA prenyltransferase domain-containing protein 1 | UBIAD1 | UbiA prenyltransferase domain-containing protein 1 OS=Homo sapiens OX=9606 GN=UBIAD1 PE=1 SV=1 | 0 | 0 | 0 | 20138000 | 13821000 | 0 | #DIV/0! | #DIV/0! | #DIV/0! |
| Q9H857 | Q9H857 | 5-nucleotidase domain-containing protein 2 | NT5DC2 | 5-nucleotidase domain-containing protein 2 OS=Homo sapiens OX=9606 GN=NT5DC2 PE=1 SV=1 | 0 | 0 | 0 | 10369000 | 17492000 | 6014300 | #DIV/0! | #DIV/0! | #DIV/0! |
| Q9NR77 | Q9NR77 | Peroxisomal membrane protein 2 | PXMP2 | Peroxisomal membrane protein 2 OS=Homo sapiens OX=9606 GN=PXMP2 PE=1 SV=3 | 0 | 0 | 0 | 13136000 | 19234000 | 0 | #DIV/0! | #DIV/0! | #DIV/0! |
| Q92581 | Q92581 | Sodium/hydrogen exchanger 6 | SLC9A6 | Sodium/hydrogen exchanger 6 OS=Homo sapiens OX=9606 GN=SLC9A6 PE=1 SV=2 | 0 | 0 | 0 | 18241000 | 14100000 | 0 | #DIV/0! | #DIV/0! | #DIV/0! |
| P14735 | P14735 | Insulin-degrading enzyme | IDE | Insulin-degrading enzyme OS=Homo sapiens OX=9606 GN=IDE PE=1 SV=4 | 0 | 0 | 0 | 15346000 | 16567000 | 0 | #DIV/0! | #DIV/0! | #DIV/0! |
| P04150;P08235 | P04150 | Glucocorticoid receptor | NR3C1 | Glucocorticoid receptor OS=Homo sapiens OX=9606 GN=NR3C1 PE=1 SV=1 | 0 | 0 | 0 | 14471000 | 12738000 | 4197100 | #DIV/0! | #DIV/0! | #DIV/0! |
| Q6PI78 | Q6PI78 | Transmembrane protein 65 | TMEM65 | Transmembrane protein 65 OS=Homo sapiens OX=9606 GN=TMEM65 PE=1 SV=2 | 0 | 0 | 0 | 16769000 | 14372000 | 0 | #DIV/0! | #DIV/0! | #DIV/0! |
| Q7L8L6 | Q7L8L6 | FAST kinase domain-containing protein 5 | FASTKD5 | FAST kinase domain-containing protein 5, mitochondrial OS=Homo sapiens OX=9606 GN=FASTKD5 PE=1 SV=1 | 0 | 0 | 0 | 15618000 | 14971000 | 0 | #DIV/0! | #DIV/0! | #DIV/0! |
| O95140 | O95140 | Mitofusin-2 | MFN2 | Mitofusin-2 OS=Homo sapiens OX=9606 GN=MFN2 PE=1 SV=3 | 0 | 0 | 0 | 13630000 | 15789000 | 0 | #DIV/0! | #DIV/0! | #DIV/0! |
| P49189 | P49189 | 4-trimethylaminobutyraldehyde dehydrogenase | ALDH9A1 | 4-trimethylaminobutyraldehyde dehydrogenase OS=Homo sapiens OX=9606 GN=ALDH9A1 PE=1 SV=3 | 0 | 0 | 0 | 8629400 | 15408000 | 5234200 | #DIV/0! | #DIV/0! | #DIV/0! |
| P82673 | P82673 | 28S ribosomal protein S35, mitochondrial | MRPS35 | 28S ribosomal protein S35, mitochondrial OS=Homo sapiens OX=9606 GN=MRPS35 PE=1 SV=1 | 0 | 0 | 0 | 0 | 19087000 | 10125000 | #DIV/0! | #DIV/0! | #DIV/0! |
| Q9Y230 | Q9Y230 | RuvB-like 2 | RUVBL2 | RuvB-like 2 OS=Homo sapiens OX=9606 GN=RUVBL2 PE=1 SV=3 | 0 | 0 | 0 | 18326000 | 0 | 10545000 | #DIV/0! | #DIV/0! | #DIV/0! |
| Q9BW92 | Q9BW92 | Threonine--tRNA ligase, mitochondrial | TARS2 | Threonine--tRNA ligase, mitochondrial OS=Homo sapiens OX=9606 GN=TARS2 PE=1 SV=1 | 0 | 16205000 | 0 | 13765000 | 16383000 | 13616000 | #DIV/0! | 1.0109843 | #DIV/0! |
| P29372 | P29372 | DNA-3-methyladenine glycosylase | MPG | DNA-3-methyladenine glycosylase OS=Homo sapiens OX=9606 GN=MPG PE=1 SV=3 | 0 | 0 | 0 | 0 | 10393000 | 15017000 | 0 | #DIV/0! | #DIV/0! |
| P31040 | P31040 | Succinate dehydrogenase [ubiquinone] flavoprotein subunit, mitochondrial | SDHA | Succinate dehydrogenase [ubiquinone] flavoprotein subunit, mitochondrial OS=Homo sapiens OX=9606 GN=SDHA PE=1 SV=2 | 0 | 0 | 0 | 18029000 | 0 | 4806400 | #DIV/0! | #DIV/0! | #DIV/0! |
| P38606 | P38606 | V-type proton ATPase catalytic subunit A | ATP6V1A | V-type proton ATPase catalytic subunit A OS=Homo sapiens OX=9606 GN=ATP6V1A PE=1 SV=2 | 0 | 0 | 0 | 0 | 11931000 | 9612600 | #DIV/0! | #DIV/0! | #DIV/0! |
| Q9H583 | Q9H583 | HEAT repeat-containing protein 1;HEAT repeat-containing protein 1, N-terminally processed | HEATR1 | HEAT repeat-containing protein 1 OS=Homo sapiens OX=9606 GN=HEATR1 PE=1 SV=3 | 5053600 | 14487000 | 0 | 178530000 | 226140000 | 21227000 | 35.327291 | 15.609857 | #DIV/0! |
| Q13488 | Q13488 | V-type proton ATPase 116 kDa subunit a isoform 3 | TCIRG1 | V-type proton ATPase 116 kDa subunit a isoform 3 OS=Homo sapiens OX=9606 GN=TCIRG1 PE=1 SV=3 | 0 | 0 | 0 | 11484000 | 9722400 | 0 | #DIV/0! | #DIV/0! | #DIV/0! |
| Q13162 | Q13162 | Peroxiredoxin-4 | PRDX4 | Peroxiredoxin-4 OS=Homo sapiens OX=9606 GN=PRDX4 PE=1 SV=1 | 0 | 0 | 0 | 16611000 | 0 | 4163800 | #DIV/0! | #DIV/0! | #DIV/0! |
| O14561 | O14561 | Acyl carrier protein, mitochondrial | NDUFAB1 | Acyl carrier protein, mitochondrial OS=Homo sapiens OX=9606 GN=NDUFAB1 PE=1 SV=3 | 0 | 0 | 0 | 7972700 | 9362300 | 1859600 | #DIV/0! | #DIV/0! | #DIV/0! |
| Q14257 | Q14257 | Reticulocalbin-2 | RCN2 | Reticulocalbin-2 OS=Homo sapiens OX=9606 GN=RCN2 PE=1 SV=1 | 0 | 0 | 65505000 | 11816000 | 5950800 | 61578000 | #DIV/0! | #DIV/0! | 0.9400504 |
| P17252;P05771;P05129 | P17252 | Protein kinase C alpha type | PRKCA | Protein kinase C alpha type OS=Homo sapiens OX=9606 GN=PRKCA PE=1 SV=4 | 30618000 | 21915000 | 0 | 48460000 | 47432000 | 16949000 | 1.5827291 | 2.1643623 | #DIV/0! |
| Q5BJH7 | Q5BJH7 | Protein YIF1B | YIF1B | Protein YIF1B OS=Homo sapiens OX=9606 GN=YIF1B PE=1 SV=1 | 11193000 | 20853000 | 0 | 84590000 | 100900000 | 16401000 | 7.5574019 | 4.8386323 | #DIV/0! |

| | | | | | | | | | | | | | |
|---|---|---|---|---|---|---|---|---|---|---|---|---|---|
| O75489 | O75489 | NADH dehydrogenase [ubiquinone] iron-sulfur protein 3, mitochondrial | NDUFS3 | NADH dehydrogenase [ubiquinone] iron-sulfur protein 3, mitochondrial OS=Homo sapiens OX=9606 GN=NDUFS3 PE=1 SV=1 | 10188000 | 0 | 0 | 14664000 | 11100000 | 5149600 | 1.4393404 | #DIV/0! | #DIV/0! |
| Q6ZRP7 | Q6ZRP7 | Sulfhydryl oxidase 2 | QSOX2 | Sulfhydryl oxidase 2 OS=Homo sapiens OX=9606 GN=QSOX2 PE=1 SV=3 | 32990000 | 30639000 | 0 | 101090000 | 95420000 | 14516000 | 3.0642619 | 3.1143314 | #DIV/0! |
| Q96RL7 | Q96RL7 | Vacuolar protein sorting-associated protein 13A | VPS13A | Vacuolar protein sorting-associated protein 13A OS=Homo sapiens OX=9606 GN=VPS13A PE=1 SV=2 | 0 | 0 | 0 | 0 | 6497100 | 7469900 | #DIV/0! | #DIV/0! | #DIV/0! |
| Q9H974 | Q9H974 | Queuine tRNA-ribosyltransferase subunit QTRTD1 | QTRTD1 | Queuine tRNA-ribosyltransferase accessory subunit 2 OS=Homo sapiens OX=9606 GN=QTRT2 PE=1 SV=1 | 6013200 | 0 | 0 | 8330500 | 8103600 | 5761100 | 1.3853689 | #DIV/0! | #DIV/0! |
| Q9Y394 | Q9Y394 | Dehydrogenase/reductase SDR family member 7 | DHRS7 | Dehydrogenase/reductase SDR family member 7 OS=Homo sapiens OX=9606 GN=DHRS7 PE=1 SV=1 | 15303000 | 32238000 | 0 | 82449000 | 104630000 | 13758000 | 5.3877671 | 3.2455487 | #DIV/0! |
| O95816 | O95816 | BAG family molecular chaperone regulator 2 | BAG2 | BAG family molecular chaperone regulator 2 OS=Homo sapiens OX=9606 GN=BAG2 PE=1 SV=1 | 0 | 0 | 0 | 0 | 10540000 | 1656500 | 0 | #DIV/0! | #DIV/0! |
| Q06124 | Q06124 | Tyrosine-protein phosphatase non-receptor type 11 | PTPN11 | Tyrosine-protein phosphatase non-receptor type 11 OS=Homo sapiens OX=9606 GN=PTPN11 PE=1 SV=3 | 14012000 | 12414000 | 0 | 36457000 | 41531000 | 11977000 | 2.6018413 | 3.345497 | #DIV/0! |
| Q96AY3 | Q96AY3 | Peptidyl-prolyl cis-trans isomerase FKBP10 | FKBP10 | Peptidyl-prolyl cis-trans isomerase FKBP10 OS=Homo sapiens OX=9606 GN=FKBP10 PE=1 SV=1 | 0 | 0 | 0 | 0 | 7651100 | 2848600 | #DIV/0! | #DIV/0! | #DIV/0! |
| P50213 | P50213 | Isocitrate dehydrogenase [NAD] subunit alpha, mitochondrial | IDH3A | Isocitrate dehydrogenase [NAD] subunit alpha, mitochondrial OS=Homo sapiens OX=9606 GN=IDH3A PE=1 SV=1 | 0 | 0 | 0 | 7362600 | 0 | 2952800 | #DIV/0! | #DIV/0! | #DIV/0! |
| Q8IYS2 | Q8IYS2 | Uncharacterized protein KIAA2013 | KIAA2013 | Uncharacterized protein KIAA2013 OS=Homo sapiens OX=9606 GN=KIAA2013 PE=1 SV=1 | 27404000 | 35920000 | 0 | 70791000 | 53711000 | 10196000 | 2.583236 | 1.4952951 | #DIV/0! |
| Q9NVH0 | Q9NVH0 | Exonuclease 3-5 domain-containing protein 2 | EXD2 | Exonuclease 3-5 domain-containing protein 2 OS=Homo sapiens OX=9606 GN=EXD2 PE=1 SV=2 | 15562000 | 0 | 0 | 11243000 | 8638700 | 1549300 | 0.722465 | #DIV/0! | #DIV/0! |
| Q9HC38 | Q9HC38 | Glyoxalase domain-containing protein 4 | GLOD4 | Glyoxalase domain-containing protein 4 OS=Homo sapiens OX=9606 GN=GLOD4 PE=1 SV=1 | 0 | 0 | 0 | 4690300 | 4998600 | 0 | #DIV/0! | #DIV/0! | #DIV/0! |
| P11142 | P11142 | Heat shock cognate 71 kDa protein | HSPA8 | Heat shock cognate 71 kDa protein OS=Homo sapiens OX=9606 GN=HSPA8 PE=1 SV=1 | 1.141E+09 | 818790000 | 327420000 | 1.205E+10 | 9.781E+09 | 4.347E+09 | 10.559839 | 11.946164 | 13.277442 |
| Q9P2E9;Q8N4C6 | Q9P2E9 | Ribosome-binding protein 1 | RRBP1 | Ribosome-binding protein 1 OS=Homo sapiens OX=9606 GN=RRBP1 PE=1 SV=5 | 19557000 | 33390000 | 101380000 | 374740000 | 390160000 | 168810000 | 19.161426 | 11.684936 | 1.6651213 |
| P11021 | P11021 | 78 kDa glucose-regulated protein | HSPA5 | Endoplasmic reticulum chaperone BiP OS=Homo sapiens OX=9606 GN=HSPA5 PE=1 SV=2 | 361250000 | 219140000 | 174890000 | 3.123E+09 | 2.679E+09 | 1.135E+09 | 8.6444291 | 12.223236 | 6.4920807 |
| Q9Y320 | Q9Y320 | Thioredoxin-related transmembrane protein 2 | TMX2 | Thioredoxin-related transmembrane protein 2 OS=Homo sapiens OX=9606 GN=TMX2 PE=1 SV=1 | 33604000 | 22397000 | 11361000 | 143970000 | 121320000 | 17471000 | 4.2843114 | 5.4167969 | 1.5378048 |
| Q9BSJ8 | Q9BSJ8 | Extended synaptotagmin-1 | ESYT1 | Extended synaptotagmin-1 OS=Homo sapiens OX=9606 GN=ESYT1 PE=1 SV=1 | 340520000 | 243160000 | 171730000 | 1.288E+09 | 1.295E+09 | 329350000 | 3.7815694 | 5.3240665 | 1.9178361 |
| P07237 | P07237 | Protein disulfide-isomerase | P4HB | Protein disulfide-isomerase OS=Homo sapiens OX=9606 GN=P4HB PE=1 SV=3 | 56255000 | 40036000 | 37883000 | 242320000 | 184950000 | 77699000 | 4.3075282 | 4.6195924 | 2.0510255 |
| Q9H3N1 | Q9H3N1 | Thioredoxin-related transmembrane protein 1 | TMX1 | Thioredoxin-related transmembrane protein 1 OS=Homo sapiens OX=9606 GN=TMX1 PE=1 SV=1 | 116220000 | 89727000 | 72616000 | 462680000 | 393220000 | 119290000 | 3.9810704 | 4.3824044 | 1.6427509 |
| Q6ZXV5 | Q6ZXV5 | Transmembrane and TPR repeat-containing protein 3 | TMTC3 | Protein O-mannosyl-transferase TMTC3 OS=Homo sapiens OX=9606 GN=TMTC3 PE=1 SV=2 | 45274000 | 25763000 | 8244200 | 129860000 | 96655000 | 24564000 | 2.8683129 | 3.7516982 | 2.9795493 |
| P63010 | P63010 | AP-2 complex subunit beta | AP2B1 | AP-2 complex subunit beta OS=Homo sapiens OX=9606 GN=AP2B1 PE=1 SV=1 | 139070000 | 121610000 | 67686000 | 403570000 | 562010000 | 101390000 | 2.9019199 | 4.6214127 | 1.4979464 |
| Q15005 | Q15005 | Signal peptidase complex subunit 2 | SPCS2 | Signal peptidase complex subunit 2 OS=Homo sapiens OX=9606 GN=SPCS2 PE=1 SV=3 | 57570000 | 49759000 | 24779000 | 212610000 | 171910000 | 43745000 | 3.6930693 | 3.4548524 | 1.7654062 |
| P04844 | P04844 | Dolichyl-diphosphooligosaccharide--protein glycosyltransferase subunit 2 | RPN2 | Dolichyl-diphosphooligosaccharide--protein glycosyltransferase subunit 2 OS=Homo sapiens OX=9606 GN=RPN2 PE=1 SV=3 | 949100000 | 702040000 | 225660000 | 3.022E+09 | 2.282E+09 | 423920000 | 3.1835423 | 3.2502422 | 1.8785784 |

| | | | | | | | | | | | | | |
|---|---|---|---|---|---|---|---|---|---|---|---|---|---|
| P21964 | P21964 | Catechol O-methyltransferase | COMT | Catechol O-methyltransferase OS=Homo sapiens OX=9606 GN=COMT PE=1 SV=2 | 379550000 | 361210000 | 129630000 | 1.194E+09 | 1.327E+09 | 175870000 | 3.1447767 | 3.6732095 | 1.3567076 |
| Q9NTJ5 | Q9NTJ5 | Phosphatidylinositide phosphatase SAC1 | SACM1L | Phosphatidylinositide phosphatase SAC1 OS=Homo sapiens OX=9606 GN=SACM1L PE=1 SV=2 | 101700000 | 85140000 | 17990000 | 269780000 | 237650000 | 48844000 | 2.652704 | 2.7912849 | 2.7150639 |
| P04843 | P04843 | Dolichyl-diphosphooligosaccharide--protein glycosyltransferase subunit 1 | RPN1 | Dolichyl-diphosphooligosaccharide--protein glycosyltransferase subunit 1 OS=Homo sapiens OX=9606 GN=RPN1 PE=1 SV=1 | 1.309E+09 | 991950000 | 522940000 | 3.784E+09 | 3.291E+09 | 819720000 | 2.8917928 | 3.3175059 | 1.5675221 |
| Q9BVK6 | Q9BVK6 | Transmembrane emp24 domain-containing protein 9 | TMED9 | Transmembrane emp24 domain-containing protein 9 OS=Homo sapiens OX=9606 GN=TMED9 PE=1 SV=2 | 117290000 | 122430000 | 46152000 | 354850000 | 312520000 | 97261000 | 3.0254071 | 2.5526423 | 2.107406 |
| O14735 | O14735 | CDP-diacylglycerol--inositol 3-phosphatidyltransferase | CDIPT | CDP-diacylglycerol--inositol 3-phosphatidyltransferase OS=Homo sapiens OX=9606 GN=CDIPT PE=1 SV=1 | 11504000 | 10793000 | 0 | 38544000 | 32689000 | 0 | 3.3504868 | 3.0287223 | #DIV/0! |
| P55060 | P55060 | Exportin-2 | CSE1L | Exportin-2 OS=Homo sapiens OX=9606 GN=CSE1L PE=1 SV=3 | 1.419E+09 | 1.557E+09 | 543460000 | 3.715E+09 | 4.661E+09 | 918990000 | 2.6189201 | 2.9935136 | 1.6909984 |
| P54136 | P54136 | Arginine--tRNA ligase, cytoplasmic | RARS | Arginine--tRNA ligase, cytoplasmic OS=Homo sapiens OX=9606 GN=RARS PE=1 SV=2 | 294350000 | 264880000 | 142930000 | 741240000 | 860180000 | 213070000 | 2.5182266 | 3.2474328 | 1.4907297 |
| Q9Y5M8 | Q9Y5M8 | Signal recognition particle receptor subunit beta | SRPRB | Signal recognition particle receptor subunit beta OS=Homo sapiens OX=9606 GN=SRPRB PE=1 SV=3 | 218720000 | 155310000 | 48008000 | 576070000 | 462850000 | 76785000 | 2.6338241 | 2.9801687 | 1.5994209 |
| P39656 | P39656 | Dolichyl-diphosphooligosaccharide--protein glycosyltransferase 48 kDa subunit | DDOST | Dolichyl-diphosphooligosaccharide--protein glycosyltransferase 48 kDa subunit OS=Homo sapiens OX=9606 GN=DDOST PE=1 SV=4 | 575120000 | 463400000 | 246330000 | 1.644E+09 | 1.378E+09 | 329710000 | 2.8581861 | 2.9730255 | 1.338489 |
| P49257 | P49257 | Protein ERGIC-53 | LMAN1 | Protein ERGIC-53 OS=Homo sapiens OX=9606 GN=LMAN1 PE=1 SV=2 | 388640000 | 237890000 | 123090000 | 886570000 | 679960000 | 222220000 | 2.2812114 | 2.8582959 | 1.8053457 |
| Q9NZJ7 | Q9NZJ7 | Mitochondrial carrier homolog 1 | MTCH1 | Mitochondrial carrier homolog 1 OS=Homo sapiens OX=9606 GN=MTCH1 PE=1 SV=1 | 92959000 | 53436000 | 8648200 | 105710000 | 122270000 | 29524000 | 1.137168 | 2.2881578 | 3.4138896 |
| Q14165 | Q14165 | Malectin | MLEC | Malectin OS=Homo sapiens OX=9606 GN=MLEC PE=1 SV=1 | 106620000 | 85392000 | 46638000 | 335910000 | 235960000 | 42936000 | 3.1505346 | 2.7632565 | 0.9206227 |
| Q9P035 | Q9P035 | Very-long-chain (3R)-3-hydroxyacyl-CoA dehydratase 3 | HACD3 | Very-long-chain (3R)-3-hydroxyacyl-CoA dehydratase 3 OS=Homo sapiens OX=9606 GN=HACD3 PE=1 SV=2 | 237420000 | 183900000 | 63514000 | 638260000 | 456290000 | 99076000 | 2.6883161 | 2.4811854 | 1.5599081 |
| Q9NZ01 | Q9NZ01 | Very-long-chain enoyl-CoA reductase | TECR | Very-long-chain enoyl-CoA reductase OS=Homo sapiens OX=9606 GN=TECR PE=1 SV=1 | 308610000 | 253150000 | 122290000 | 872370000 | 580210000 | 190480000 | 2.8267717 | 2.2919613 | 1.557609 |
| Q92544 | Q92544 | Transmembrane 9 superfamily member 4 | TM9SF4 | Transmembrane 9 superfamily member 4 OS=Homo sapiens OX=9606 GN=TM9SF4 PE=1 SV=2 | 102830000 | 67149000 | 15991000 | 224730000 | 187300000 | 26081000 | 2.1854517 | 2.7893193 | 1.6309799 |
| Q00610;P53675 | Q00610 | Clathrin heavy chain 1 | CLTC | Clathrin heavy chain 1 OS=Homo sapiens OX=9606 GN=CLTC PE=1 SV=5 | 2.485E+09 | 2.461E+09 | 804740000 | 5.664E+09 | 6.555E+09 | 1.27E+09 | 2.2796201 | 2.6634025 | 1.5776524 |
| P16435 | P16435 | NADPH--cytochrome P450 reductase | POR | NADPH--cytochrome P450 reductase OS=Homo sapiens OX=9606 GN=POR PE=1 SV=2 | 133000000 | 80562000 | 35566000 | 283010000 | 177580000 | 71748000 | 2.1278947 | 2.204265 | 2.0173199 |
| O95292 | O95292 | Vesicle-associated membrane protein-associated protein B/C | VAPB | Vesicle-associated membrane protein-associated protein B/C OS=Homo sapiens OX=9606 GN=VAPB PE=1 SV=3 | 377460000 | 255420000 | 149110000 | 1.015E+09 | 607770000 | 181880000 | 2.6892916 | 2.3794926 | 1.2197706 |
| P30101 | P30101 | Protein disulfide-isomerase A3 | PDIA3 | Protein disulfide-isomerase A3 OS=Homo sapiens OX=9606 GN=PDIA3 PE=1 SV=4 | 184330000 | 139450000 | 79843000 | 410950000 | 346910000 | 123080000 | 2.2294255 | 2.4877017 | 1.5415252 |
| Q7Z434 | Q7Z434 | Mitochondrial antiviral-signaling protein | MAVS | Mitochondrial antiviral-signaling protein OS=Homo sapiens OX=9606 GN=MAVS PE=1 SV=2 | 23465000 | 30180000 | 10137000 | 62584000 | 61613000 | 13790000 | 2.6671212 | 2.0415176 | 1.360363 |
| Q8N5K1 | Q8N5K1 | CDGSH iron-sulfur domain-containing protein 2 | CISD2 | CDGSH iron-sulfur domain-containing protein 2 OS=Homo sapiens OX=9606 GN=CISD2 PE=1 SV=1 | 66510000 | 51003000 | 19907000 | 152240000 | 110930000 | 27943000 | 2.2889791 | 2.1749701 | 1.4036771 |
| P00387 | P00387 | NADH-cytochrome b5 reductase 3;NADH-cytochrome b5 reductase 3 membrane-bound form;NADH-cytochrome b5 reductase 3 soluble form | CYB5R3 | NADH-cytochrome b5 reductase 3 OS=Homo sapiens OX=9606 GN=CYB5R3 PE=1 SV=3 | 909620000 | 782580000 | 240520000 | 2.024E+09 | 1.823E+09 | 312630000 | 2.2247752 | 2.3299854 | 1.2998087 |

| | | | | | | | | | | | | | |
|---|---|---|---|---|---|---|---|---|---|---|---|---|---|
| O95573 | O95573 | Long-chain-fatty-acid--CoA ligase 3 | ACSL3 | Long-chain-fatty-acid--CoA ligase 3 OS=Homo sapiens OX=9606 GN=ACSL3 PE=1 SV=3 | 457400000 | 404000000 | 142120000 | 980760000 | 827880000 | 228350000 | 2.1442064 | 2.0492079 | 1.6067408 |
| Q07812 | Q07812 | Apoptosis regulator BAX | BAX | Apoptosis regulator BAX OS=Homo sapiens OX=9606 GN=BAX PE=1 SV=1 | 45544000 | 41561000 | 18488000 | 96077000 | 98533000 | 21035000 | 2.1095424 | 2.3708044 | 1.137765 |
